# Distinct super-enhancer elements differentially control *Il2ra* gene expression in a cell-type specific fashion

**DOI:** 10.1101/2022.11.18.516445

**Authors:** Rosanne Spolski, Peng Li, Vivek Chandra, Boyoung Shin, Chengyu Liu, Jangsuk Oh, Min Ren, Yutaka Enomoto, Erin E. West, Stephen Christensen, Edwin C.K. Wan, Meili Ge, Jian-Xin Lin, Pandurangan Vijayanand, Ellen V. Rothenberg, Warren J. Leonard

## Abstract

The IL-2 receptor α-chain (IL-2Rα/CD25) is constitutively expressed on DN2/DN3 thymocytes and Treg cells but induced by IL-2 on mature T and NK cells. *Il2ra* expression is regulated by a super-enhancer extensively bound by STAT5 in mature T cells. Here, we demonstrate that STAT5 cooperates with Notch to induce/maintain *Il2ra/*CD25 expression in DN2/DN3 cells. Moreover, we systematically investigated CD25 regulation using a series of mice with deletions spanning STAT5 binding elements. Deleting the upstream super-enhancer region mainly affected constitutive CD25 expression on DN2/DN3 thymocytes and Tregs, whereas deleting an intronic region primarily decreased IL-2-induced CD25 on peripheral T and NK cells. Thus, distinct elements preferentially control constitutive versus inducible expression in a cell-type-specific manner, with the MED1 coactivator co-localizing with specific STAT5 binding sites. Moreover, the intronic region was a dominant element whose deletion altered the structure throughout the super-enhancer in mature T cells. These results demonstrate differential functions for distinct super-enhancer elements, thereby indicating ways to manipulate CD25 expression in a cell-type specific fashion.

## Introduction

Super-enhancers are extended regions of chromatin that bind transcription factors and coactivators at high density and orchestrate the formation of nuclear condensates, driving transcription of key genes involved in lineage establishment or maintenance (Hnisz et al., 2013; Whyte et al., 2013). Super-enhancers are variably considered to represent distinctive structures or to be comprised of multiple individual typical enhancers (Hay et al., 2016; Hnisz *et al*., 2013; Pott and Lieb, 2015), and they are dynamic structures that can be remodeled during differentiation or in response to external signals (Adam et al., 2015), with context-dependent roles. Coactivators such as MED1 and BRD4 can form phase-separated condensates at super-enhancers, thereby concentrating the transcription apparatus at key cell-identity genes and influencing their expression. Moreover, studies of super-enhancers have provided insights into the mechanisms underlying gene control in normal and pathologic states (Chong et al., 2018; Sabari et al., 2018). Super-enhancer activity can be modulated by 3-dimensional chromatin interactions organized into compartments denoted as topologically-associating domains (TADs) (Rowley and Corces, 2018).

The human and mouse genes encoding IL-2Rα (CD25) have been extensively studied over the years to explain the responsiveness of this gene to both activation via the T cell receptor and IL-2. Our lab and others identified a series of upstream positive regulatory regions (PRRs), including PRRI, PRRII, PRRIII, where PRRI and PRRII were required for the mitogenic induction of the *IL2RA* gene and PRRIII was an IL-2 response element, as well as a CD28 response element (Cross et al., 1987; Cross et al., 1989; John et al., 1995; John et al., 1996; Kim et al., 2001; Kim and Leonard, 2002; Rameil et al., 2000; Soldaini et al., 1995; Yeh et al., 2001; Yeh et al., 2002). PRRIII bound STAT5A and STAT5B proteins as well as ELF-1, HMG-I(Y), and a GATA-1-like protein. Our lab then characterized another IL-2-regulated element, PRRIV, that could bind STAT5 proteins and HMG-I(Y) (Kim et al., 2006; Liao et al., 2013). We subsequently showed that the *Il2ra* super-enhancer is the top-ranked IL-2/STAT5-dependent super-enhancer in mouse pre-activated T cells, comprising approximately 13 STAT5 binding sites and exhibiting potent activity upon IL-2 stimulation (Li et al., 2017). The corresponding super-enhancer in the human *IL2RA* locus has a similar overall organization (Li *et al*., 2017). We previously deleted three individual STAT5 binding sites (one upstream and two intronic) in mice using CRISPR/Cas9 technology and showed that each deletion partially contributed to IL-2-induced *Il2ra* expression in mature CD8^+^ T cells. Moreover, the mutation in mice of an autoimmunity risk variant in an intronic enhancer delayed but did not abrogate *Il2ra* gene expression upon TCR stimulation in mature T cells (Simeonov et al., 2017). However, it is not clear how individual super-enhancer elements throughout the gene contribute to cell-type specific *Il2ra* expression during development, in different lineages, and in response to stimuli.

To further understand *Il2ra* regulation, here we have generated a range of individual or combinatory deletions of STAT5-bound regulatory elements in the *Il2ra* super-enhancer *in vivo* and analyzed them in multiple cell types, including in response to cytokine stimulation. CD25 expression varies at different stages of mouse lymphoid development. It is constitutively expressed in subsets of double negative (DN) thymocytes (DN2/DN3) and in regulatory T cells (Treg cells); in contrast, CD25 is not expressed in resting T or NK cells but is induced in response to appropriate antigen or cytokine signals (Leonard et al., 2019; Liao *et al*., 2013). We show that STAT5 is important for normal CD25 expression at specific DN stages of T cell development as well as in mature CD8^+^ T cells. We reasoned that focusing on the STAT5 binding sites would allow us to identify important enhancer elements that regulated expression during development or in response to T cell receptor or cytokine signals. Strikingly, we found that distinct super-enhancer regions preferentially control constitutive versus inducible CD25 expression at different stages of T cell development and in other cell types as well. Furthermore, we show extensive looping within this super-enhancer, with intronic regions dominantly affecting the overall chromatin structure and activity of the gene, and moreover, we show the co-localization of the MED1 coactivator and of transcription factors NFATc1, FOXP3, TCF1, and SMAD4 at the super-enhancer. This detailed analysis provides mechanistic insights into CD25 regulation not only by IL-2 but also by TCR and TGFβ signals throughout early T cell development as well as in specific functional T cell subsets.

## Results

### Lineage- and developmental-specific STAT5 binding and open chromatin structure at the *Il2ra* locus

We initially examined the STAT5 binding profile at the *Il2ra* locus in multiple immune cell populations using ChIP-Seq (**Figure 1A**; the top enriched STAT5-binding GAS (gamma-activated sequence) motifs in each cell type are shown in **Table S1**, and the GAS motif at each upstream and intronic site is shown in **Table S2)**. TCR-activated CD8^+^ T cells and inducible Treg (iTreg) cells had similar STAT5 binding profiles in response to IL-2 stimulation, with most of the upstream and intronic sites occupied, whereas little STAT5 binding was observed in these cells in the absence of IL-2 stimulation (**Figure 1A**). Distinctive STAT5 binding patterns were found in natural Treg cells isolated ex vivo, as well as in natural killer (NK) cells stimulated with IL-15, splenic dendritic cells (DCs) stimulated with GM-CSF, and in mast cells stimulated with anti-IgE (**Figure 1A**), suggesting that distinct *Il2ra* super-enhancer elements might be differentially utilized in a cell-type-specific fashion.

**Figure 1.**
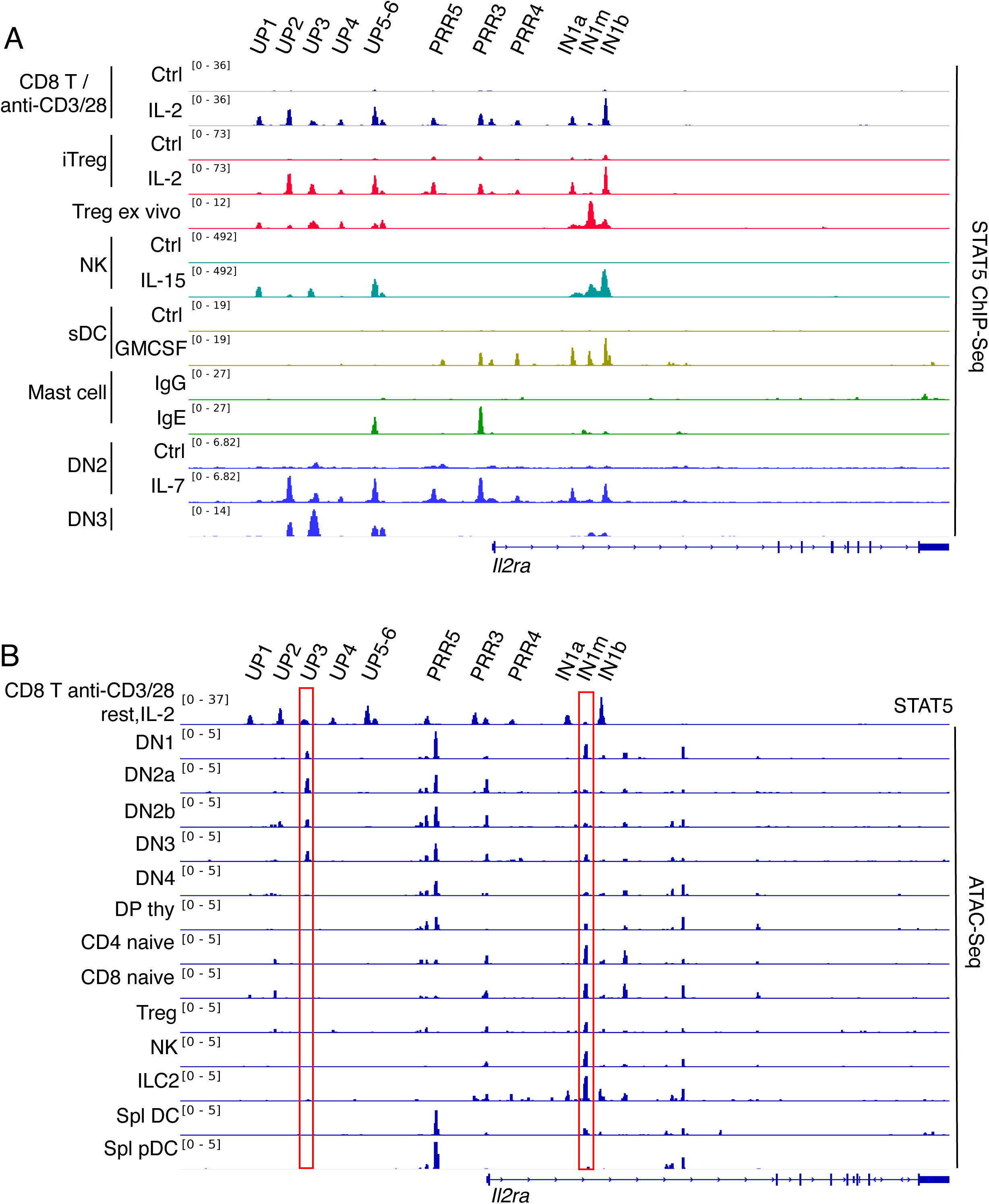
Lineage and developmental chromatin structure at the *Il2ra* super-enhancer. **(A)** STAT5 binding as assessed by ChIP-Seq experiments using *in vitro* activated lymphoid (CD8^+^ T cells, iTreg cells, NK cells, and DN2 thymocytes) and myeloid populations (splenic DCs and mast cells) or *ex vivo* populations (DN3 and Treg cells). The CD8^+^ T cells were pre-activated with anti-CD3 + CD28 for 2 days, rested for 4 h, and then not stimulated or stimulated with IL-2 for 4 h. All ChIP-Seq experiments were performed on two independent samples, with similar results. Representative data are shown. **(B)** Open chromatin regions in the *Il2ra* locus as assessed by ATAC-Seq. Red rectangles highlight regions that appear to be developmentally significant. Data are from the Immgen database.

During early mouse T cell development, CD25 is a key marker of DN thymocytes that is expressed in the DN2 (cKit^+^CD44^+^CD25^+^) and DN3 (cKit^−^CD44^−^CD25^+^) stages bridging T-cell lineage commitment (Rothenberg et al., 2008). These cells upregulate IL-7Rα as they enter the DN2 stage, and the natural thymic microenvironmental signal, IL-7, stimulates these cells, activating STAT5A and/or STAT5B. Although *Stat5a* and *Stat5b* deletion severely compromises early T cell development (Yao et al., 2006) and in the individual absence of either STAT5A or STAT5B, there is a partial defect in T cell numbers and/or IL-2-induced CD25 expression (Imada et al., 1998; Nakajima et al., 1997), whether CD25 expression in DN thymocytes is dependent on STAT5 activation and binding to the *Il2ra* gene has remained unclear. To investigate this, we first identified the stage at which STAT5 is activated by IL-7 by using B6.Bcl11b-mCitrine reporter mice (**Figure S1A**) and found that STAT5 phosphorylation approximately corresponded to when CD25 is first expressed in DN thymocytes—namely in DN2a/DN2b cells and then extending into DN3 cells but declining in DN4 cells (**Figures S1B-S1D**). Next, we used sgRNAs to acutely delete both *Stat5a* and *Stat5b* in normal Cas9;*Bcl2*-transgenic hematopoietic precursors, starting one stage before CD25 is normally expressed, and we examined the early stages of their T-lineage development in artificial thymic organoid (ATO) (**Figures S1E-S1G**) and OP9-DLL1 coculture (**Figures S1H-S1J**) in vitro systems. The presence of a *Bcl2* transgene in each system prevented the diminished survival that otherwise would have resulted from defective IL-7-STAT5 mediated induction of *Bcl2* in these DN cells. In the ATO system, deletion of either *Stat5a* or *Stat5b* partially lowered pSTAT5 in DN2 cells, and DN2 cells generated from *Stat5a*/*Stat5b*-deleted precursors showed greatly diminished IL-7-induced pSTAT5 (**Figures S1F and S1G**). To measure the impact of STAT5 loss on CD25 expression, we generated DN2 cells using the OP9-DLL1 co-culture T-cell differentiation system (Romero-Wolf et al., 2020; Shin et al., 2021) (**Figure S1H**) and found that *Stat5a*/*Stat5b*-deletion reduced CD25 expression intensity across the stages (**Figure S1I**). Importantly, this effect on CD25 expression did not reflect a developmental block given that *Stat5a/Stat5b*-deleted cells still underwent normally timed commitment to the T-cell lineage, as shown by upregulation of a *Bcl11b*-mCherry reporter allele. By day 7, the control cells had progressed to the DN2 and early DN3 stages, with strong surface expression of CD25; however, when both *Stat5a* and *Stat5b* were deleted, both surface CD25 (**Figures S1I and S1J)** and *Il2ra* RNA **(Figures S1K and S1L)** expression were significantly reduced in both pre-commitment (BCL11b^-^) and post-commitment (BCL11b^+^) DN2 stages. A global transcriptomic analysis showed that *Stat5a/Stat5b-*deficient DN2/3 cells had lower expression of growth and viability control genes (e.g., *Bcl2, Bcl2l1, Socs2, Cdkn1a, Eno1, Xbp1,* and *Bhlhe40*), as well as of *Il2ra* (**Table S3**), but the cells were not altered in their developmental progression (**Table S3)** as defined by a curated panel of markers, including *Tcf7, Bcl11b, Lck, Thy1, Cd3* clusters*, Ets1*, *Tcf12,* and *Rag* genes (Zhou et al., 2019). This effect of STAT5 was in addition to the known requirement for Notch signaling to induce and sustain CD25 expression in DN thymocytes (**Figure S1M;** this panel is from (Romero-Wolf *et al*., 2020)). Thus, STAT5 activation cooperated with Notch signaling to induce and maintain normal levels of *Il2ra/*CD25 expression in DN2-DN3 pro-T cells.

In vitro generated DN2 (CD25^+^cKit^+^) thymocytes responded to IL-7 with a STAT5 binding pattern similar to that observed in IL-2-stimulated CD8^+^ T cells (**Figure 1A**), but interestingly, steady-state thymic DN3 cells isolated *ex vivo* had their greatest STAT5 binding at the upstream region of the super-enhancer, primarily at the UP3 element (**Figure 1A**), a pattern distinct from that observed in IL-2-induced CD8^+^ T cells or iTregs. Other factors including Notch/RBPJ (Chen et al., 2019; Del Real and Rothenberg, 2013; Liu et al., 2010a; Romero-Wolf *et al*., 2020), RUNX1 (Guo et al., 2008; Shin *et al*., 2021), BCL11b (Liu et al., 2010b), and TCF-1 (Weber et al., 2011) are known to contribute to lineage progression to the DN2b-DN3 stages, and these factors bound in proximity to the STAT5 binding sites in DN2b-DN3 cells **(Figure S2**), suggesting that they might cooperate to control the developmental regulation of *Il2ra* expression in DN cells.

We next analyzed chromatin accessibility at the *Il2ra* locus throughout lymphoid development using ATAC-Seq (Assay for Transposase-Accessible Chromatin with high-throughput sequencing) datasets from the ImmGen database (Yoshida et al., 2019). Interestingly, an open chromatin region at upstream element UP3 (**Figure 1B**; left red box) was detected in DN1 thymocytes, modestly increased in DN2a, DN2b, and DN3 thymocytes, but then was essentially absent in DN4 thymocytes; thus, the loss of chromatin accessibility at UP3 approximately correlated with the loss of CD25 expression. Accessibility at the intronic IN1m STAT5 binding site was relatively low in thymocyte populations, was greater in naïve CD4 and CD8 cells, Treg cells, NK cells, and ILC2s, but then was weak in splenic conventional DC and plasmacytoid DC (pDC) populations (**Figure 1B**, right red box), suggesting differential use of upstream and intronic elements in different cellular populations.

### Upstream super-enhancer elements are required for CD25 expression in DN thymocytes

To investigate the roles of the upstream and intronic enhancer elements in lymphoid development and function, we next deleted a range of individual elements or combinations of elements within the *Il2ra* super-enhancer (**Figure 2A**) *in vivo* using CRISPR-Cas9 methodology. No individual upstream element deletion nor even deletion of the entire UP1-6 upstream region (ΔUP1-6) or intron region (ΔIntron) or both together (ΔUP1-6/ΔIntron) significantly affected either thymic or splenic cellularity **(Figures S3A and S3B),** nor did they affect the percentage of DN thymocytes **(Figure S3C)**. However, mice lacking the UP1-6 region had a profound decrease in the percentage of CD25^+^CD44^+^ DN2 and CD25^+^CD44^-^ DN3 thymocytes, whereas loss of the intron region had little effect (**Figure 2B**). RNA-Seq analysis of DN thymocytes from WT, ΔUP1-6, or ΔUP1-6/ΔIntron mutant mice showed that *Il2ra* mRNA was essentially absent in the mutant mice, but *Bcl11b, Tcf7,* and *Il7ra* mRNAs, which are normally expressed in DN2/DN3 cells (Hosokawa and Rothenberg, 2018), were still expressed **(Figure S3D)**, indicating that pro-T cells were indeed still present but lacked CD25 expression due to deletion of essential *Il2ra* regulatory elements. Separate deletions of UP1, UP2, UP3, and UP5-6 did not affect CD25 expression in DN thymocytes (**Figures 2C and 2D**). Interestingly, however, deletion of the ΔUP2-3 region, which extends downstream of UP3 to include a putative RBPJ binding site (TGACTAATG) (Romero-Wolf *et al*., 2020), essentially eliminated CD25 expression on DN cells. We therefore generated mice lacking UP3 extending through the RBPJ site (ΔUP3-RBPJ) and found that this deletion also abrogated CD25 expression on DN thymocytes, phenocopying the ΔUP1-6 mice (**Figures 2C and 2D**). Thus, the UP3-RBPJ region was indispensable for *Il2ra*/CD25 expression in DN thymocytes. To further analyze the role of the upstream elements, we cloned individual upstream enhancer elements in a reporter construct and analyzed reporter activity in a DN3-like cell line (SCID-ADH-2C2) (Anderson et al., 2002; Dionne et al., 2005). The UP3-RBPJ region was sufficient to drive reporter activity, but reporter activity was not augmented by stimulating the cells with IL-2, IL-7, or the combination of IL-2 with PMA + ionomycin (PI), which can mimic TCR stimulation (**Figure 2E**). Strikingly however, activity was significantly reduced when the STAT5 binding site (GAS motif), the RBPJ binding site, or both sites were mutated (**Figure 2F**), or when a Notch inhibitor (γ-secretase inhibitor, GSI) was used (**Figure 2G**). Even though *in vivo* deletion of the UP5-UP6 region had little if any effect, the UP5 element also exhibited modest reporter activity in these cells, with enhanced activity induced by IL-2, IL-7, PI, and PI/IL-2 (**Figure 2E**), but inhibiting Notch had little effect on its activity, indicating the specificity of the Notch inhibitor for the UP3-RBPJ region (**Figure 2G**). These results indicate a key role for the UP3-RBPJ element in regulating CD25 expression in DN2 and DN3 cells, with its activity requiring both STAT5 and Notch. In addition to the co-localization of RBPJ with STAT5 at the UP3 site, RUNX1 and BCL11b, which are both involved in progression through DN thymocyte development, also bound at the UP3 region **(Figure S2)**, suggesting that multiple signals may be involved in the regulation of CD25 expression in DN2/DN3 thymocytes.

**Figure 2.**
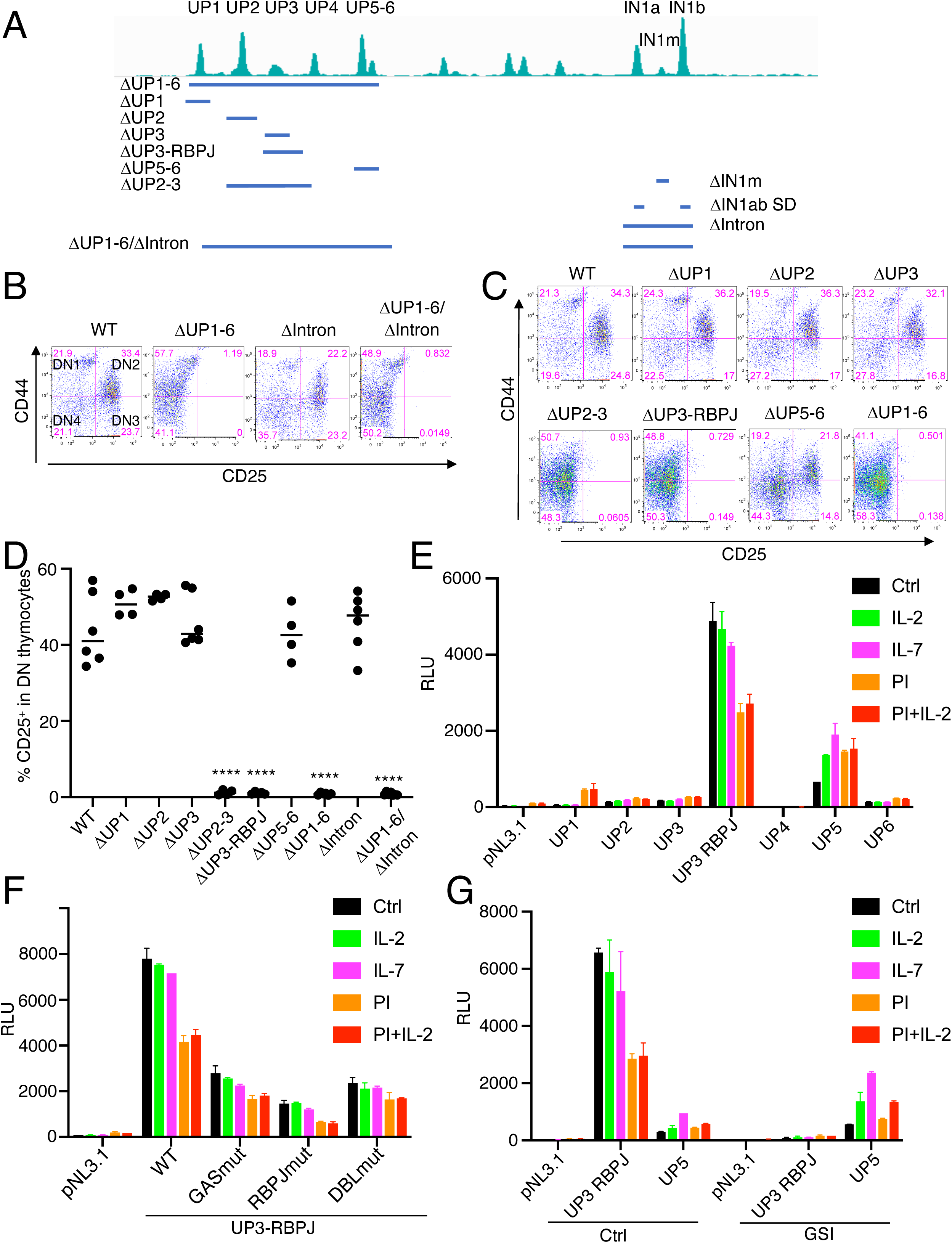
Upstream enhancer elements control constitutive CD25 expression in DN thymocytes. **(A)** Schematic summary of CRISPR-Cas9 generated mutant mice. Indicated are the upstream or intronic regions that were deleted; these mutants were used to determine the functional importance of individual or groups of elements within the *Il2ra* super-enhancer. **(B)** Flow cytometry profiles gated on CD4^-^CD8^-^ double negative thymocytes from mice containing either the UP1-6 upstream deletion (ΔUP1-6), the intron deletion (Δintron), or deletion of both regions. The data are representative of 3 experiments. **(C and D)** Representative flow cytometry data of DN thymocytes from the indicated mutant mice, with data from multiple (two to three) experiments summarized in **(D)**. Shown is the percentage of DN cells that express CD25. **(E)** *In vitro* transfection of a DN3 cell line (SCID-ADH), with individual upstream (UP) regions cloned into pNL3.1 reporter construct. **(F)** A WT UP3/RBPJ reporter construct or reporters containing mutation of the GAS motif, the RBPJ motif, or both motifs were transfected into the DN3 cell line. **(G)** Treatment of the cells with a γ-secretase inhibitor to block Notch signaling reduced reporter expression by the UP3/RBPJ construct but did not block expression from a UP5 reporter construct. E-G are representative of 3 experiments.

### Differential control of CD25 expression on thymic and peripheral Treg cells

Because CD25 is also constitutively expressed on Treg cells, we next investigated whether *Il2ra* gene regulation in these cells was similar to what we observed in DN thymocytes. Thymic and splenic Treg cell numbers were not significantly affected by any of the deletions **(Figures S4A and S4B)** but deleting UP1-6 or both UP1-6 and intronic regions markedly reduced CD25 expression on both thymic and splenic Tregs (**Figures 3A and 3B**). Strikingly, whereas the ΔUP2-3 or ΔUP3-RBPJ deletions essentially abrogated CD25 expression on DN2/DN3 thymocytes (**Figure 2C**), these deletions did not significantly affect CD25 expression on thymic Tregs (**Figure 3A**) and only modestly lowered CD25 expression on splenic Tregs (**Figure 3B**), indicating distinctive regulation of the *Il2ra* gene in DN thymocytes and Tregs. Although the intron deletion did not significantly affect CD25 expression on thymic DN2/DN3 cells (**Figure 2B**), ΔIntron thymic FoxP3^+^ Tregs include a particularly strong CD25-negative population (**Figure 3C** left and middle panels **and 3D)**, reminiscent of FoxP3^+^CD25^-^ thymic Treg precursor cells that normally express CD25 in response to IL-2, IL-7, or IL-15 (Cheng et al., 2013; Owen et al., 2018). This suggests that cytokine-mediated induction of CD25 in precursor Tregs may be particularly dependent on the intronic region, although a population of CD25^low^ thymocytes was also present in the ΔUP1-6 mice (**Figure 3C**). Consistent with this, when the total thymic CD4^+^ cells were examined for Treg precursor populations, it was apparent that deletion of either the intron or UP1-6 region led to a decreased percentage of CD25^+^FoxP3^neg^ cells but increased percentage of CD25^neg^FoxP3^+^ cells **(Figures S4C and S4D)**. This CD25^neg^FoxP3^+^ precursor population was increased in ΔIntron and ΔUP1-6/ΔIntron Tregs (**Figures 3D** **and S4B)**. In ΔΙntron splenic Tregs, such a bimodal distribution was not as readily observed (**Figure 3C**, right panel), indicating differences in CD25 regulation in thymic and splenic Tregs and suggesting that CD25^neg^FoxP3^+^ Tregs in the thymus either do not exit to the periphery or are eliminated.

**Figure 3.**
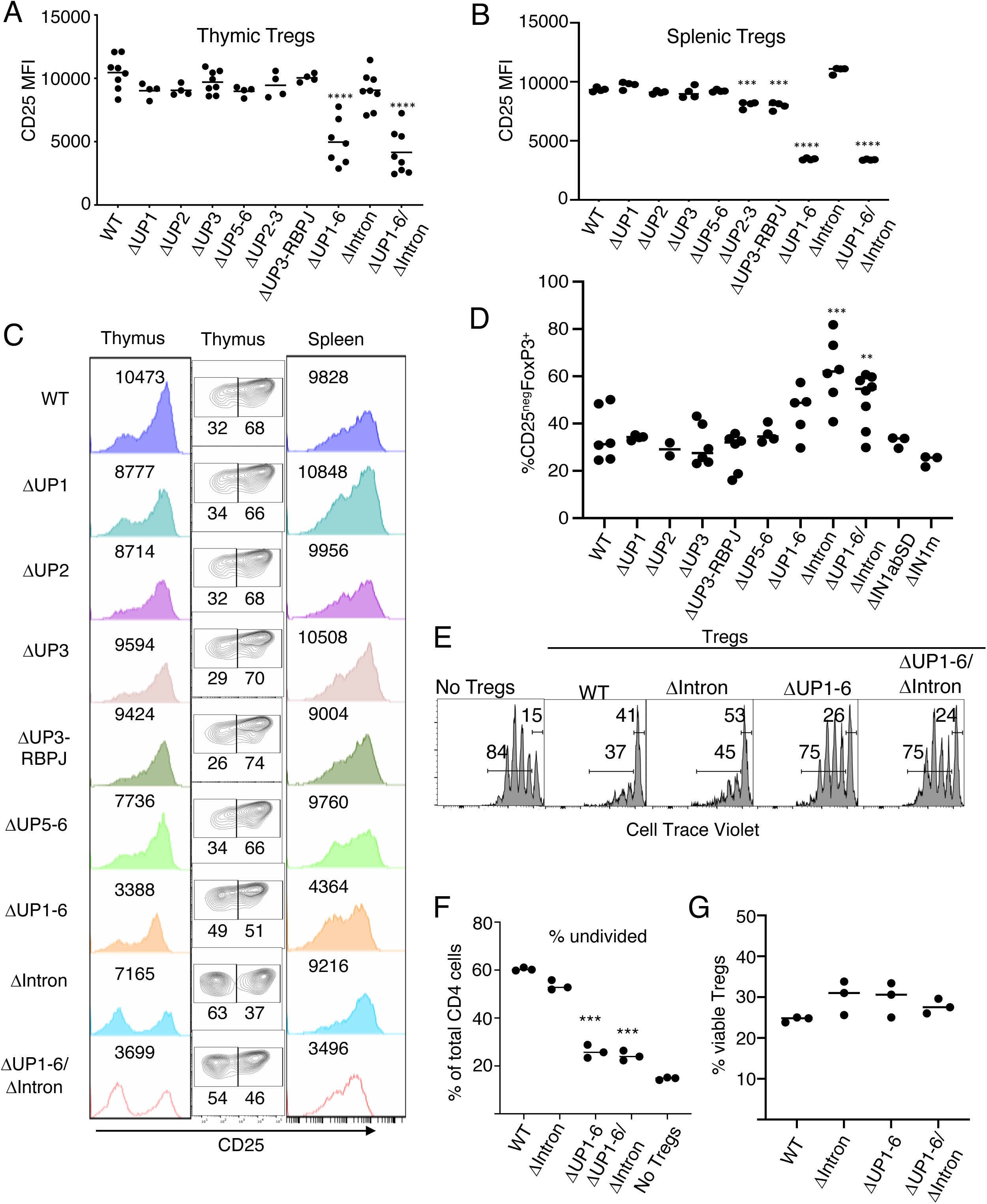
Upstream enhancer elements control the level of constitutive CD25 expression in both thymic and splenic Treg cells. **(A-C)** Flow cytometry profiles for CD25 expression (MFI) gated on CD4^+^FoxP3^+^ thymic and splenic cells from mice with upstream and intronic deletions in the *Il2ra* super-enhancer. Data for thymic Tregs (**A**) and splenic Tregs (**B**) were combined from two to three experiments with one representative experiment in **(C)** showing the histograms of thymic Tregs on the left and splenic Tregs on the right panel. The middle panel gates on the CD25^low^ and CD25^hi^ thymic Tregs. (**D**) Summary (from 2-3 experiments) of the percentage of CD25^low^ cells within the thymic FoxP3^+^ population of mice with deletions of upstream or intronic enhancer regions. **(E and F)** Splenic Treg suppressive activity as measured by inhibition of WT CD4^+^ T cell proliferation (see Methods). Cell Trace Violet dilution at day 3 CD4^+^ T cells co-cultured with purified splenic Tregs from WT, ΔIntron, ΔUP1-6 or ΔUP1-6/ΔIntron mice; shown are a representative experiment (**E**) and the % undivided cells from 3 mice (**F**). (**G**) Viability of Tregs at the end of the suppression assay, as assessed by flow cytometry.

Interestingly, ΔIntron mice expressed lower levels of FoxP3 in thymic Tregs, while ΔUP1-6 mice expressed lower levels of FoxP3 mainly in splenic Tregs **(Figure S4E)**. This suggests that STAT5-activating cytokines such as IL-2 can regulate expression of FoxP3 in Tregs, consistent with evidence that a FoxP3 enhancer element containing STAT5 binding sites can mediate IL-2-regulated Treg function (Dikiy et al., 2021). In addition to the regulation of FoxP3 transcription by IL-2-induced STAT5 activation, FoxP3 binding sites colocalized with the STAT5 sites in the *Il2ra* intronic and proximal promoter regions (**Figure S5**), suggesting a feedback regulation where IL-2 via STAT5 regulates expression of FoxP3, which in turn regulates *Il2ra* expression and thus responsiveness to IL-2 in the Treg lineage. Splenic Tregs with the ΔUP1-6 deletion had lower CD25 than WT Tregs or those with the ΔIntron deletion (**Figure 3C**) and less potently suppressed CD4^+^ T cell proliferation in vitro, as compared to WT or ΔΙntron Tregs (**Figures 3E and 3F**), importantly correlating reduced CD25 levels and reduced levels of FoxP3 with impaired function in the Treg lineage. Tregs from ΔUP1-6 mice had similar viability to WT and ΔΙntron Tregs at the end of the suppression assay (**Figure 3G**), excluding death as the reason for reduced suppression.

### Inducible CD25 expression on CD8^+^ and CD4^+^ T cells is controlled by super-enhancer intronic elements

We next assessed the effects of upstream and intron deletions in the super-enhancer on inducible CD25 expression in peripheral CD8^+^ T cells. In freshly isolated WT CD8^+^ T cells, IL-2 induced *Il2ra* mRNA expression, with greater induction at 1000 U/ml than 100 U/ml IL-2 (**Figure S6A**). Cells from ΔUP1-6 mice exhibited similar induction with 100 U IL-2, but for unclear reasons, unlike WT cells, the response was reproducibly not further increased with 1000 U IL-2. Strikingly, the intron deletion abolished IL-2-induced *Il2ra* mRNA expression **(Figure S6A)**. We next investigated the roles of the individual upstream or intronic elements in cell surface CD25 expression in response to IL-2, either in the absence or presence of anti-CD3 + anti-CD28 stimulation. IL-2-induced CD25 expression at day 2 of culture generally paralleled *Il2ra* mRNA expression, with much lower CD25 expression on the ΔIntron or ΔUP1-6/ΔIntron cells (**Figure 4A**, left 3 sets of flow profiles, and **Figure 4B**). When CD8^+^ T cells were stimulated with anti-CD3 + anti-CD28 and then IL-2, CD25 expression was much higher (**Figure 4A**, right 3 sets of flow profiles, and **Figure 4B**), but deleting the intron essentially eliminated CD25 expression (**Figure 4B**). None of the individual upstream deletions decreased CD25 expression as much as the larger ΔUP1-6 deletion (**Figures 4A** and **4B**), suggesting that elements within this region cooperatively regulate CD25 expression in CD8^+^ T cells, but even ΔUP1-6 did not affect expression nearly as much as deleting the intronic region. Thus, the intron region contains the major elements responsive to TCR and IL-2 signals in CD8^+^ T cells, including a TCR-responsive intronic element approximately 200 bp from the 3’ end of IN1a (Simeonov *et al*., 2017) as well as TCR-responsive NFATc1 sites that colocalize with each of the intronic STAT5 sites (**Figure S5**) all of which are deleted in the ΔIntron deletion, although the upstream region also contributes to CD25 inducibility, particularly at 4 days (**Figures 4C** **and S6B**). Deletion of both IN1a and IN1b or of IN1m **(Figure S6C)** or of the element described by Simeonov (Simeonov *et al*., 2017) had more modest defects than deletion of the entire intron region, suggesting that the intronic elements functionally cooperate to mediate high level IL-2-induced *Il2ra* expression in CD8^+^ T cells.

**Figure 4.**
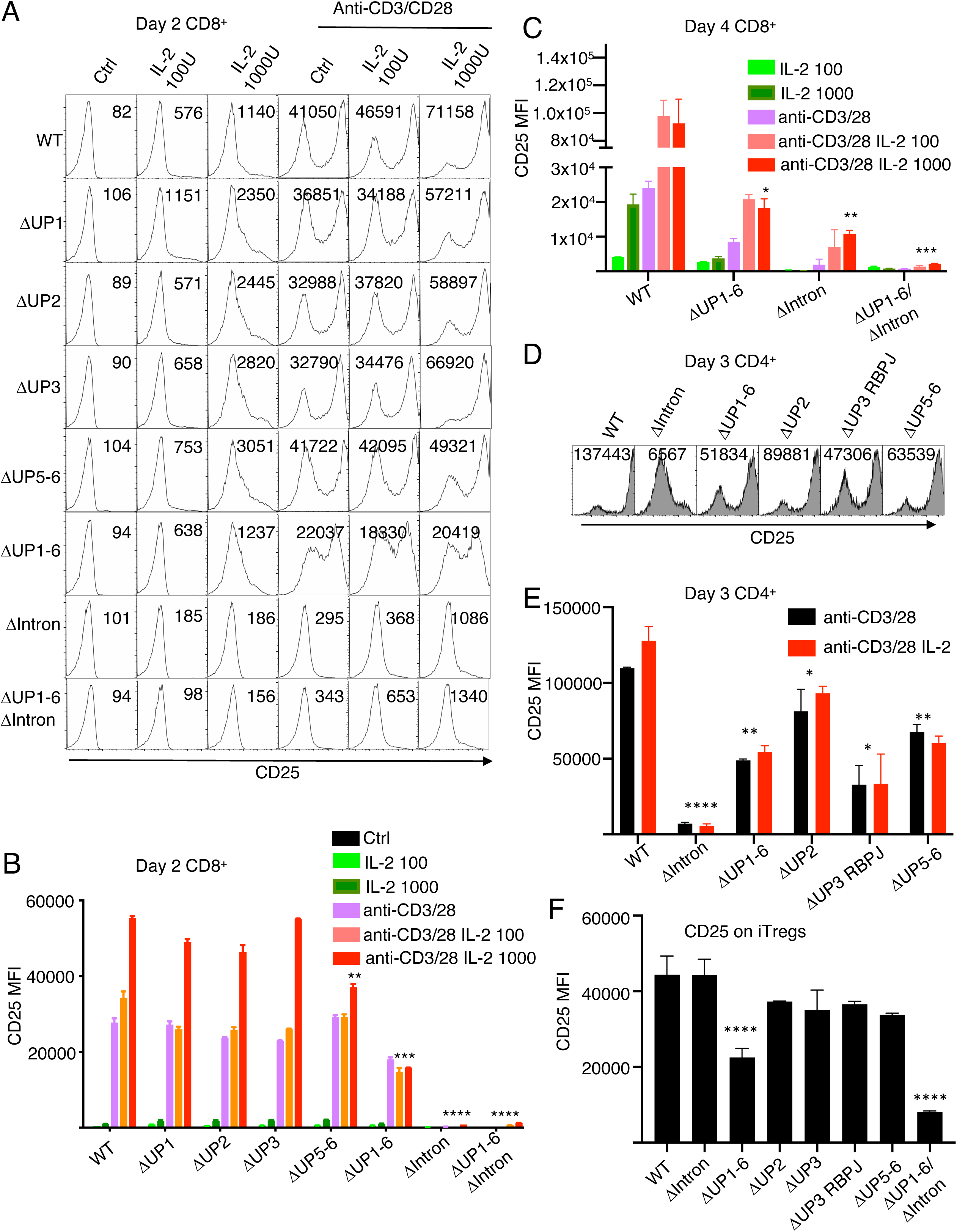
Inducible CD25 expression in CD8^+^ and CD4^+^ T cells is strongly regulated by intronic enhancer elements. **(A and B)** Flow cytometric measurement of CD25 surface expression on CD8^+^ T cells isolated from WT or the indicated mutant mice and stimulated with or without IL-2 in the presence or absence of anti-CD3 + anti-CD28. Representative histograms at day 2 are shown in **(A)** with collective MFI data from 2 experiments depicted in **(B)**. (**C**) Summary of flow cytometric measurement of CD25 surface expression on CD8^+^ T cells isolated from WT or the indicated mutant mice and stimulated with or without IL-2 in the absence or presence of anti-CD3 + anti-CD28. Analysis was performed at day 4. **(D-E)** CD4^+^ T cells isolated from WT or the indicated mutant mice and from the indicated mice were activated with anti-CD3 + anti-CD28 in the presence of IL-2 for 3 days, and CD25 expression was measured by flow cytometry. Representative profiles are shown in **(D)**, and collective profiles are shown in **(E)**. **(F)** Naïve CD4^+^ T cells from the indicated mice were activated under iTreg conditions with TGFβ + IL-2 + anti-IFNγ and anti-IL-4 for 3 days. CD25 expression was then measured by flow cytometry. **A-F** are representative of 3 independent experiments.

Because naïve CD4^+^ T cells do not express IL-2Rα and have low expression of IL-2Rβ, they cannot substantially respond to IL-2 prior to activation (Smith et al., 2017). We therefore analyzed the effects of *Il2ra* upstream and intron deletions on the CD4^+^ T cell response to anti-CD3 + anti-CD28, which augments expression of IL-2Rα and IL-2Rβ. In contrast to the IL-2-induced CD25 expression in CD8^+^ T cells co-stimulated with anti-CD3 + anti-CD28 (**Figures 4A-4C**), in WT CD4^+^ T cells, CD25 was strongly expressed after TCR stimulation, with no further increase induced by IL-2 (**Figures 4D** and **4E**). This was presumably at least in part due to endogenous IL-2 production induced by TCR signaling in CD4^+^ T cells. Flow cytometric analysis revealed an essential role for the intron and a partial role for UP1-6 for the induction of CD25 in CD4^+^ T cells (**Figures 4D and 4E**), similar to their roles in CD8^+^ T cells. As compared to WT, individual deletion of UP2, UP3-RBPJ and UP5-6 each had partial effects on CD25 expression in CD4^+^ T cells (**Figure 4E**). However, deleting IN1a and IN1b together or IN1m had essentially no effect, in contrast to the effect of deleting the combined region (ΔIntron) **(Figure S6D)**, suggesting cooperative effects.

As noted above, splenic Tregs as a whole exhibited distinctive regulatory requirements as compared to those of thymic Tregs. Interestingly, for iTregs derived from the splenic CD4^+^ population, ΔIntron had no effect, ΔUP1-6 significantly reduced CD25 expression, and loss of both regions further reduced CD25 expression (**Figure 4F**), indicating key differences in the elements used in iTregs versus peripheral T cells. None of the individual upstream elements was solely responsible for the lower CD25 expression in iTregs observed with ΔUP1-6 (**Figure 4F**). Because iTregs are induced in the presence of TGFβ, we examined WT CD4^+^ T cells activated with anti-CD3 + anti-CD28 in the presence of IL-2, TGFβ, or both TGFβ and IL-2. Compared to anti-CD3 + anti-CD28, the addition of IL-2 had little effect, and although the addition of TGFβ repressed CD25, combining IL-2 with TGFβ reversed this repression, restoring CD25 expression to the level observed with anti-CD3 + anti-CD28 **(Figure S7A)**. Similar qualitative effects were observed with cells from ΔUP1-6 mice, but interestingly, CD4^+^ and CD8^+^ T cells from ΔIntron mice had minimal responses to IL-2, whereas the combination of TGFβ with IL-2 had a strong cooperative effect on CD25 expression on ΔIntron cells **(Figures S7A and S7B)**. Because of the effect of TGFβ, we reasoned that SMAD4 might bind to this region. We previously identified a TGFβ responsive element in the upstream region proximal to the promoter (in a region that we denoted positive regulatory region 5, PRR5) (Kim et al., 2005), and when we examined published ChIP-Seq data for SMAD4, we found SMAD4 binding sites throughout the *Il2ra* locus (Zhang et al., 2017) **(Figure S5)**, co-localizing with STAT5 binding sites in the intron and others, including at PRR5. These latter SMAD4 sites potentially operate independently of the intronic region and might help to explain the difference in intron-dependence of CD25 expression in CD4^+^ T cells versus iTregs. These data collectively indicate that multiple signals converge at distinct regions of this gene and can serve to differentially modulate CD25 expression in different lineages.

### *Il2ra* super-enhancer intronic elements control cytokine induced CD25 expression on NK cells

NK cells potently respond to IL-15 as well as IL-2, with IL-15 and STAT5 being critical for the development and proliferation of these cells (Di Santo, 2006; Lin et al., 2017; Yao *et al*., 2006). Because IL-12 can augment CD25 expression in NK cells both in vivo and in vitro (Lee et al., 2012), we examined CD25 induction in splenic NK cells cultured with IL-15 alone or with IL-12 and found cooperative induction (see **Figure 5A** for day 4 and **Figure 5B** for days 2 and 4). As anticipated, IL-15-stimulated NK cells expressed *Il2ra* mRNA (**Figure 5C**), with higher levels induced by IL-15 + IL-12, although even this level of expression (**Figure 5B**) was still significantly lower than the level induced by IL-2 on WT CD8^+^ T cells based on a comparison of MFIs (**Figure 4C**). As compared to WT NK cells, NK cells from the ΔIntron and ΔIntron/ΔUP1-6 mutant mice had markedly decreased CD25 expression in response to IL-15 + IL-12, whereas ΔUP1-6 NK cells were intermediate in their response at both the protein (**Figure 5B**) and mRNA (**Figure 5C**) levels.

**Figure 5.**
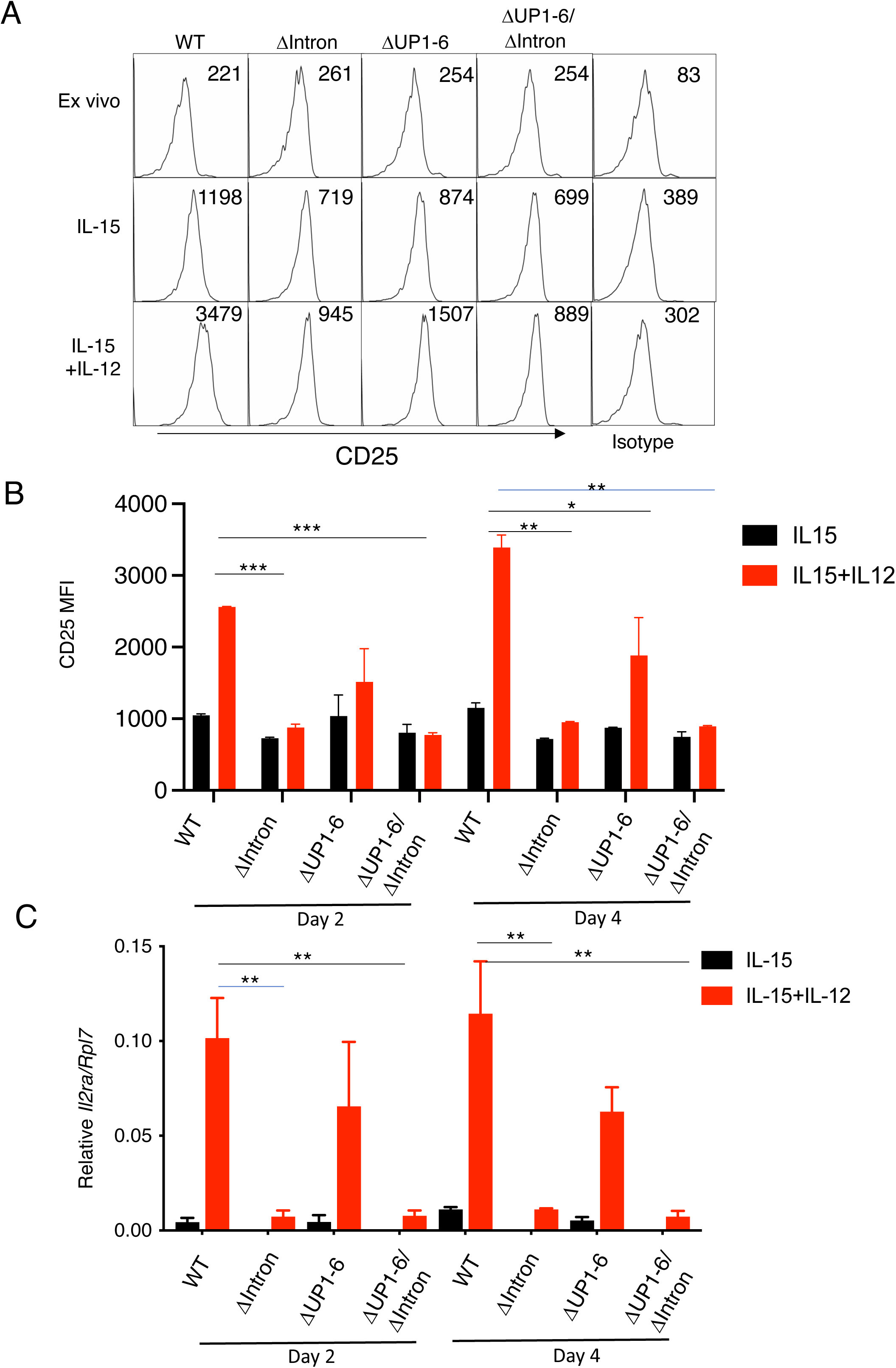
Cytokine regulation of CD25 expression in NK cells is controlled by intronic elements. **(A)** NK cells were purified from WT, Δintron, ΔUP1-6, or Δintron/ΔUP1-6 mice, CD25 was measured by flow cytometry *ex vivo* and then after 4 days of culture with either IL-15 or IL-15 + IL-12. **(B)** NK cells were stimulated in vitro for 2 days (left panel) or 4 days (right panel) with IL-15 or IL-15 + IL-12, and flow cytometry was used to measure CD25 expression. **(C)** RT-PCR was used to measure *Il2ra* mRNA relative to *Rpl7*. In panel **C**, for those samples where the IL-15 signal is not visible, it was not separable from the baseline. **A-C** are representative of two independent experiments.

### Intronic elements control the chromatin structure of the *Il2ra* super-enhancer

Above, the ATAC-Seq data showed that the intronic IN1m region had an open chromatin structure throughout lymphoid development, and UP3-RBPJ was accessible in DN2 and DN3 but not in DN4 thymocytes (**Figure 1B**), correlating with CD25 expression. Histone-3-lysine27-acetylation (H3K27Ac), which is a mark of active enhancers (Creyghton et al., 2010), was evident in DN3 thymocytes from *Rag2* KO mice and in DN thymocytes from WT mice and was most enriched at UP3-RBPJ and promoter regions, whereas H3K27Ac was lower at the intronic region (green tracks, top of **Figure 6A**). Deleting the intronic region (ΔIntron) in DN thymocytes did not substantially affect H3K27Ac in the upstream region or at the promoter (**Figure 6A**), consistent with the upstream rather than intronic region controlling constitutive CD25 expression on DN2-DN3 thymocytes (**Figure 2B**). In WT CD8^+^ T cells, H3K27Ac was distributed throughout the *Il2ra* super-enhancer region. In contrast to the results in DN2-DN3 thymocytes, deleting the intron region in CD8^+^ T cells reduced H3K27Ac levels throughout the *Il2ra* locus, including at the upstream region, whereas deleting UP1-6 in CD8^+^ T cells resulted in a loss of H3K27Ac signal in the upstream region as expected but had little if any effect in the intronic region (**Figure 6A**). As expected, the changes in chromatin structure were specific to the *Il2ra* locus; for example, the various *Il2ra* deletions did not affect the H3K27Ac binding profile at the locus for another IL-2-induced gene, *Cish* **(Figure S8A)**.

**Figure 6.**
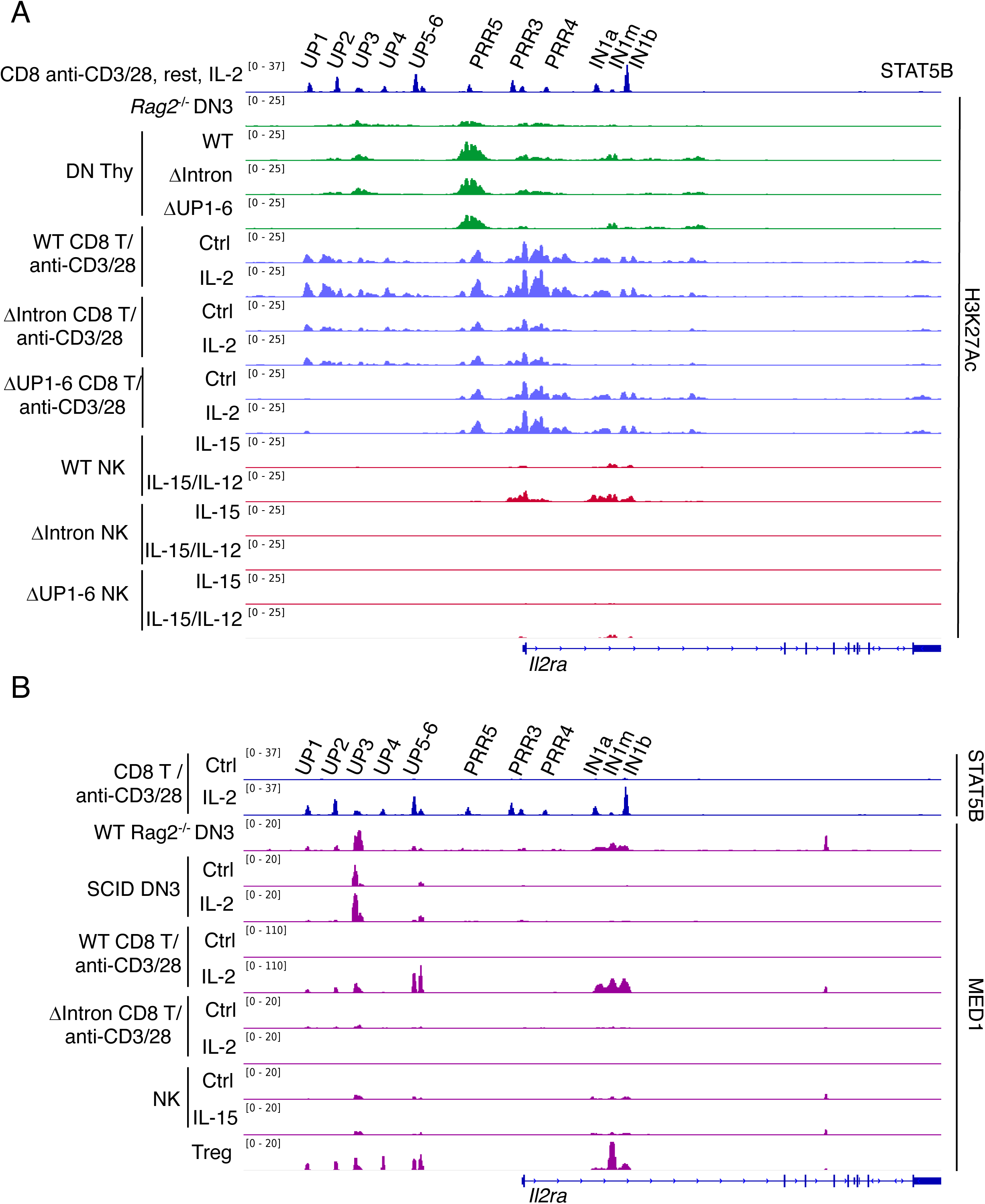
Super-enhancer structure is distinct at different developmental stages. **(A)** ChIP-Seq profiles for H3K27Ac marks were assessed in *ex vivo* isolated *Rag2*^-/-^ DN3 thymocytes, DN thymocytes from WT, ΔIntron and ΔUP1-6 mice; in WT, ΔIntron and ΔUP1-6 CD8^+^ T cells that were stimulated in vitro with anti-CD3 + anti-CD28 without or with 1000 U IL-2 for 48 hrs; and in NK cells that were stimulated with IL-15 or IL-15 + IL-12. **(B)** ChIP-Seq profiles for MED1 were assayed in DN3 thymic cells, in vitro activated CD8^+^ T cells and NK cells, and *ex vivo* Treg cells. All ChIP-Seq results are representative of two independent experiments. All ChIP-Seq experiments in **A** and **B** were performed on two independent samples, with similar results. Representative data are shown.

We also evaluated H3K27Ac marks at the *Il2ra* locus in WT, ΔIntron, and ΔUP1-6 NK cells stimulated with IL-15 or IL-15 + IL-12 and found that they were primarily in the promoter and intronic regions and not evident in the upstream region (bottom six tracks, **Figure 6A**). H3K27Ac binding intensity at the *Il2ra* locus was enhanced in NK cells stimulated with IL-15 + IL-12 as compared to stimulation with IL-15 alone (**Figure 6A**), consistent with augmented protein and mRNA expression (**Figures 5A-5C**). This enhancement was also associated with greater numbers of total STAT5 binding sites identified by ChIP-Seq analysis in NK cells stimulated with IL-15 + IL-12 versus IL-15 alone (**Figures S8B and S8C**). Interestingly, deleting either the UP1-6 region or the intron led to the loss of H3K27Ac marks throughout the gene (**Figure 6A**), indicating the importance of both regions for controlling *Il2ra* expression in NK cells. Consistent with the observation that *Il2ra* gene expression is significantly lower in NK cells than in CD8^+^ T cells or CD4^+^ T cells, an analysis of STAT5-bound super-enhancers in CD8^+^ and NK cells showed that the super-enhancer score of the gene was ranked first in CD8^+^ T cells but was only 151^st^ in NK cells **(Figure S9).**

These observations demonstrate that the enhancer activity and chromatin structure at the *Il2ra* gene are dynamic, with substantial variation during lymphoid development and in different cell types, suggesting a potential hierarchical chromatin organization within this super-enhancer. The intronic element influences the chromatin interactions of the locus and expression in CD8^+^ T cells and NK cells, whereas the upstream super-enhancer elements seemed to exhibit broad effects but were most dramatically required for *Il2ra* expression in thymic DN2 and DN3 cells with lesser effects in CD8^+^ T cells, consistent with normal *Il2ra* expression in DN thymocytes from ΔIntron mice. Note that for CD8^+^ T cells, TCR activation results in strong H3K27Ac, with little subsequent effect of IL-2. This is consistent with our previous observations that IL-2 augments STAT5 binding but has no evident further effect on H3K27Ac (Li *et al*., 2017).

### Recruitment of MED1 by the *Il2ra* super-enhancer

Strong binding of MED1, a core component of the Mediator co-activator that is required for the activation of RNA polymerase II, is characteristic of super-enhancers (Whyte *et al*., 2013). We therefore analyzed the profile of MED1 binding to the *Il2ra* super-enhancer in various cell types. In DN3 thymocytes and in a DN3 cell line, which constitutively express CD25 and have strong STAT5 binding at UP3 with closely juxtaposed RBPJ binding (**Figure 1**), there was significant MED1 binding to the UP3 element (**Figure 6B**). In WT CD8^+^ T cells stimulated with IL-2, MED1 partially co-localized with STAT5B binding in both the upstream and intronic regions (**Figure 6B**). Genome-wide analysis of STAT5B and MED1 binding in CD8^+^ T cells revealed that 44% of genes bound by STAT5 also had co-binding of MED1 **(Figure S10A)**, suggesting possible cooperation between these factors. We examined published RNA-Seq data from preactivated CD8^+^ T cells that were stimulated with IL-2 for 48 hours (Hermans et al., 2020) and found that 25% of the 100 most-inducible genes were co-bound by STAT5 and MED1, including *Cish* and *Il2ra* **(Figure S10B)**. We next examined the binding of the cohesin complex component RAD21, given the ability of RAD21 and MED1 to form a complex, connecting gene expression and chromatin architecture (Kagey et al., 2010). Indeed, consistent with the binding pattern observed for MED1 (**Figure 6B**), there was significant binding of RAD21 at the UP3 and intronic regions in wild-type CD8^+^ T cells (**Figure S10C)**.

Interestingly, ΔIntron CD8^+^ T cells had essentially no MED1 binding throughout the *Il2ra* locus (**Figure 6B**), consistent with the loss of H3K27Ac marks (**Figure 6A**) and suggesting that deleting the intron disrupted the super-enhancer structure in these cells. Moreover, there was only weak MED1 binding in NK cells cultured with or without IL-15 (**Figure 6B**), consistent with the lower CD25 protein expression on IL-15-stimulated NK cells (**Figure 5B**) than on IL-2-stimulated CD8^+^ T cells (**Figure 4C**). Treg cells constitutively express CD25 and had a MED1 binding profile relatively similar to that observed in IL-2-induced CD8^+^ T cells (**Figure 6B**). These results collectively underscore the differential use of elements within the super-enhancer during lymphoid development and/or cytokine stimulation.

To further investigate the role of the higher order chromatin structure at the *Il2ra* locus, we performed H3K27Ac HiChIP experiments using CD8^+^ T cells stimulated with anti-CD3 + anti-CD28. These experiments revealed significant H3K27Ac-mediated chromatin interactions among the *Il2ra* promoter, the upstream and the intronic cis-regulatory elements, with increased looping in the presence of IL-2 (**Figure 7A**) in WT cells. Interestingly, there was also looping to the adjacent *Il15ra* gene, although RNA-Seq showed no significant difference *in Il15ra* expression in WT, ΔIntron, and ΔUP1-6 mice. The ΔIntron cells not only lacked all loops from within the intron region (as expected) but had greatly reduced looping from the upstream region to the *Il2ra* promoter (**Figure 7A**), which in part may reflect the lower H3K27Ac (**Figure 6A**). Conversely, ΔUP1-6 eliminated all loops from within the upstream region, but it had relatively little effect on looping from the intron to the promoter (**Figure 7A**). Other IL-2-induced gene loci showed no difference in looping (e.g., the *Bcl2* locus; **Figure 7B**), confirming that the changes in chromatin interactions were specific to the *Il2ra* locus. These data underscore important contributions of both the intronic and upstream super-enhancer regions to the regulation of the *Il2ra* gene in mature T cells.

**Figure 7.**
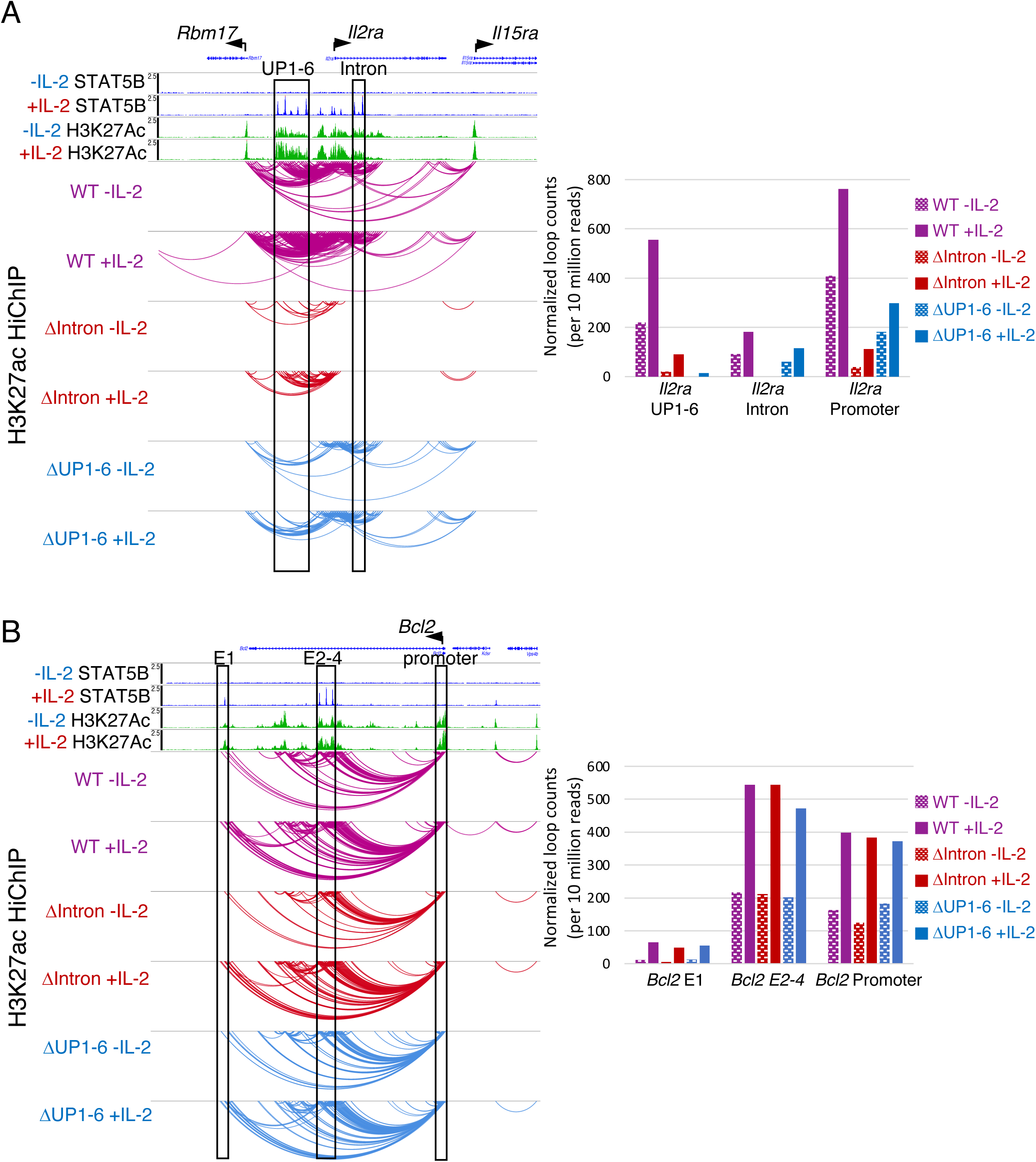
Higher order chromatin looping within the *Il2ra* super-enhancer is disrupted by the loss of the intronic region. H3K27Ac HiChIP assay showing chromatin interactions at the *Il2ra* **(A)** and *Bcl2* **(B)** loci in CD8^+^ T cells from WT, ΔIntron, or ΔUP1-6 mice that were activated with anti-CD3 + anti-CD28 and then not stimulated or stimulated with IL-2. H3K27Ac HiChIP results are representative of two independent experiments.

## Discussion

In this study, we have extensively dissected the *Il2ra* super-enhancer *in vivo* and identified regulatory elements that differentially control gene expression in distinct cellular lineages and at different stages of lymphoid development, as well as in response to different cytokines. Strikingly, the upstream region controls constitutive CD25 expression during early thymic development as well as in Treg cells, but it also contributes to expression in all lymphoid cell types studied, whereas the intronic region is primarily vital for inducible CD25 expression in mature T cells. The distinctive biological roles for different super-enhancer elements in the *Il2ra* locus contrasts to findings in the globin super-enhancer where different elements were found to exhibit largely additive effects.

In the *Il2ra* upstream region, we identified a Notch-responsive element that specifically controls constitutive DN2/DN3 thymocyte CD25 expression and cooperates with STAT5, thus linking the critical Notch signaling pathway to a STAT5-regulated transcriptional event. In this regard, the UP3 region has an open chromatin configuration during early thymic development, suggesting that this region is primed to respond to the combination of STAT5 and Notch signals. In the DN2/DN3 thymocytes, the importance of UP3-RBPJ is consistent with the indispensability of NOTCH/RBPJ input (Romero-Wolf *et al*., 2020), and it is possible that binding of RUNX1, BCL11b, and TCF1 to some of the upstream elements is also critical. Notch signaling is important for T cell commitment in the thymic microenvironment, and the acquisition of CD25 expression is thus linked to these developmentally important signals (Adler et al., 2003; Radtke et al., 2013).

Unlike the upstream super-enhancer region, the intronic region was most critical for IL-2-inducible *Il2ra* expression in mature CD8^+^ T cells and in NK cells, and selective individual deletion of the three STAT5 binding sites within this region had only minor effects on *Il2ra* mRNA expression, indicating that the elements exhibit cooperative effects for full expression. Moreover, deletion of the intron disrupted chromatin structure throughout the *Il2ra* locus, for example, as measured by reduced MED1 and H3K27Ac binding. In contrast to this effect on inducible expression in mature T cells, deleting the intron did not affect CD25 expression on DN2/DN3 thymocytes, indicating that the structure of the super-enhancer may evolve as cells traverse through development to mature T cells. Interestingly, although the intron deletion did not affect CD25 expression on peripheral Tregs, it altered the distribution of developing FoxP3^+^ cells in the thymus, such that CD25^neg^FoxP3^+^ precursors accumulated, suggesting that the intronic region can also control the cytokine-inducible expression of CD25 in this precursor population.

In this study, we analyzed the role of STAT5 in progression through early thymic development, demonstrating a key requirement for this transcription factor for the expression of CD25 in the earliest DN stages. Moreover, we demonstrated co-localization of STAT5 with other transcription factors (NOTCH, RUNX, BCl11b, TCF1) that facilitate development. It is likely that some of these factors individually or in combination can associate with other nuclear proteins that orchestrate chromatin accessibility and that such pioneer factors then allow cytokine-dependent recruitment of STAT5 to these sites. This situation may be analogous to that in developing Treg cells where FOXP3 binds to previously established enhancers to orchestrate the functional transcriptional program of Tregs (Samstein et al., 2012). Indeed, lower CD25 expression on Treg cells correlates with a less suppressive phenotype, and FOXP3 binds to both intronic and promoter *Il2ra* STAT5 site regions, suggesting that FOXP3 may contribute to the induction of the *Il2ra* gene and thereby to a positive feedback loop.

Context-dependent regulation of genes ranging from *even-skipped* in Drosophila to *Myc* in mammals has long been known to be mediated by tissue-specific activities of distinct enhancers, sometimes widely separated in the genome (Bahr et al., 2018; Fujioka et al., 1999; Herranz et al., 2014; Kieffer-Kwon et al., 2013; Small et al., 1993). However, super-enhancers are usually identified by the simultaneous activity of distinct cis-regulatory elements within the same cellular context. Here, we show that the *Il2ra* super-enhancer comprises both modes of regulation. Some super-enhancers have individual enhancer elements that function in an additive fashion (Hay *et al*., 2016; Shin et al., 2016), while others possess hierarchical structures in which some individual elements are dominant in their influence on the rest of the super-enhancer (Huang et al., 2018). Our data indicate that the intronic element of the *Il2ra* super-enhancer functions as such a hub in mature T cells, with increased occupancy of STAT5 binding sites and enhanced histone modifications associated with active chromatin, consistent with global effects on the entire locus when the intron is deleted. MED1 co-localized with STAT5 protein at three sites within the intron, and this association was induced by cytokine stimulation. Moreover, MED1 binding correlated with the transcriptional activity of the *Il2ra* locus, with lower binding in NK cells than in mature CD8^+^ T cells. In addition, cohesin also co-localized at STAT5 binding sites in the intron and at the UP3 element, perhaps indicative of co-occupancy of these factors at super-enhancer condensates. STAT proteins have been shown to be incorporated into nuclear condensates following cytokine stimulation (Zamudio et al., 2019). Cohesin and MED1 have been shown to function as coactivators in a complex of proteins that bind with lineage-specific or inducible transcription factors at the enhancers of active genes, leading to the distinctive architecture of chromatin proteins that define the super-enhancer structure (Kagey *et al*., 2010; Zamudio *et al*., 2019). In CD8^+^ T cells, the binding of MED1 and cohesin was greatly diminished throughout the super-enhancer by the loss of the intronic hub, suggesting that this locus may be displaced from the super-enhancer condensate.

Overall, our findings underscore that the *Il2ra* super-enhancer, which is the highest ranked STAT5-based super-enhancer in both IL-2-induced CD4^+^ and CD8^+^ T cells (Li *et al*., 2017), critically regulates gene expression, with subregions that differentially influence expression of this critical gene in distinct cellular populations. Our observations therefore clarify the basis for differential regulation of the *Il2ra* gene in distinct cell types. More broadly, they illustrate how distinct elements within a super-enhancer can collectively control constitutive versus inducible gene expression in different cell types. In the lineage choices confronting T cells, the interaction of cytokine induced STAT5 activation with pre-existing chromatin structures at distinct stages of development leads to plasticity in *Il2ra* regulation, as has been observed in other lineages (Adam *et al*., 2015).

Although the general organization of the super-enhancer is similar in mouse and human in that it has both upstream and intronic elements, most elements are not rigorously conserved between these species (Li *et al*., 2017). There are many autoimmune-related SNPs at the *Il2ra* gene locus (Farh et al., 2015), but so far, only one common human SNP associated with protection from type 1 diabetes has been studied. This SNP led to delayed TCR-induced CD25 expression in T cells but did not affect steady state CD25 levels on Tregs, and the mechanism for protection from type I diabetes remains unclear (Simeonov *et al*., 2017). Interestingly, the *Il2ra* elements that are bound by STAT5 differ across cell types; for example, there are variations in occupancy of upstream versus downstream elements in CD8^+^ T cells versus Treg cells or DCs. Accordingly, selective interference with binding at key positions can be predicted to differentially affect CD25 expression in a cell type-related fashion, with presumed parallels in the human and potentially in human disease. For example, intronic mutations/deletions can be predicted to affect CD25 expression on NK cells and potentially impact innate immunity, whereas upstream mutations/deletions might instead impact Treg function and thus autoimmunity, an area for future investigation. Our study also provides a vision of how one might differentially target and specifically control expression of the *Il2ra* gene in different populations of cells. Specifically, the differential roles of upstream elements (e.g., for Treg cells) versus intronic elements (e.g., for effector CD8^+^ T cells or NK cells) might potentially allow one to differentially influence expression of CD25 on distinct cellular populations, with potential therapeutic utility.

## Acknowledgments

This work was supported by the Division of Intramural Research, National Heart, Lung, and Blood Institutes, NHLBI (W.J.L); NIH grants R01AI135200 and R01HD100039 (E.V.R.); R24AI108564, R01AI121426 and R01HL114093 (V.P.). B.S. was supported by a Cancer Research Institute-Irvington Postdoctoral Fellowship. For ChIP-Seq and RNA-Seq analysis, DNA sequencing was performed in the NHLBI DNA Sequencing Core. We thank Dr. Ning Du for help with mast cell STAT5 ChIP-Seq studies.

## Author Contributions

R.S. and P.L. designed the project, performed experiments, interpreted data, and wrote the manuscript. V.C. and P.V performed HiChIP experiments and analyzed the data. P.L and S.C. performed computational analysis. B.S. and E.V.R. performed the DN thymocyte in vitro experiments and ChIP-Seq experiments and analyzed data and wrote the manuscript. C.L. and J.O. designed and constructed mouse deletion mutants. M.R, Y.E, E.E.W., E.W., J.O., M.G., and J.L. contributed to in vitro cloning, animal experiments, and ChIP-Seq experiments. W.J.L. supervised the project and wrote the manuscript.

## Accession numbers

For the RNA expression in control and *Stat5a;Stat5b* double knockout DN2 cells, the GSE accession number is GSE184845 (https://www.ncbi.nlm.nih.gov/geo/query/acc.cgi?acc=GSE184845).

For the RNA-Seq data from IL-2-induced CD8^+^ T cells (Hermans *et al*., 2020), the GSE accession number is GSE143903. Other data can be viewed at https://nih.box.com/s/6bmco09put8ilgo1pgqd2xfdfzrrmff9 and raw data will be deposited in GEO database prior to publication.

## Materials and Methods

### Mice

CRISPR-Cas9 mutant mice were generated by the transgenic core of the National Heart, Lung, and Blood Institute (NHLBI), as previously described (Li *et al*., 2017). All experiments at NHLBI were performed using protocols approved by the NHLBI Animal Care and Use Committee and followed NIH guidelines for use of animals in intramural research. sgRNAs used to generate the mutant mice and the sequence coordinates of deletions are in **Table S3**. For in vitro derived DN2-DN3 cells with acute *Stat5a; Stat5b* deletion, mice combining a *ROSA26-Cas9* knock-in and a *Bcl2* transgene were bred at the California Institute of Technology from B6.Cg-Tg(BCL2)25Wehi/J(Bcl2-tg) and B6.Gt(ROSA)26Sortm1.1(CAG-cas9*,-EGFP)Fezh/J (Cas9) mice, originally purchased from the Jackson Laboratory. When a Bcl11b reporter allele was required, these mice were further bred with B6. *Bcl11b^mCh/mCh^* (mCherry) reporter mice, as previously described (Shin *et al*., 2021). All protocols followed NIH Guidelines and were approved by the California Institute of Technology Animal Care and Use Committee.

### Cell isolation and culture

CD4^+^, CD8^+^ and Tregs were isolated using StemCell kits. NK cells were purified from spleens using a Miltenyi kit and expanded in IL-15 (10 ng/ml) for 10 days for use in ChIP-Seq experiments. For short term analysis of NK cells, purified NK cells were incubated with IL-15 (10 ng/ml) in the presence or absence of IL-12 (10 ng/ml) for 2-4 days. CD8^+^ T cells were cultured in RPMI-1640 medium containing 10% FBS and stimulated with IL-2 (100 or 1000 U/ml) or anti-CD3/anti-CD28 for 2 days, rested for 4 hours and then IL-2 added for the indicated times. For iTreg cultures, naïve CD4^+^ T cells were isolated from spleens and cultured with TGFβ (2 ng/ml), IL-2 (1000 U/ml), anti-IFNγ (10 μg/ml), and anti-IL-4 (10 μg/ml) for 4 days. DN thymocytes were enriched by depletion of total thymus with anti-CD4 and anti-CD8 antibodies from Miltenyi. Enriched DN3 cells were isolated from *Rag2* KO mice. Splenic DCs were purified using a Miltenyi kit, rested for 1 h, and stimulated with GM-CSF (20 ng/ml) for 2 h. Bone marrow-derived mast cells were generated by culturing bone marrow from femurs for 4-6 weeks in medium containing 10 ng/ml stem cell factor and 10 ng/ml IL-3; cells were sensitized with 0.5 μg/ml IgE (557079, Sigma) overnight, and stimulated with 50 ng/ml DNP-HSA (human serum albumin)(A6661, Sigma). For DN2 and DN3 cell culture, bone marrow progenitor cells from 8-12 week-old mice with a C57BL/6J background were first obtained by depleting mature cells expressing CD3LJ (clone 145-2C11), CD19 (clone 1D3), B220 (clone RA3-6B2), NK1.1 (clone PK136), CD11b (clone M1/70), CD11c (clone N418), Ly6G/C (clone RB6-8C5), and Ter119 (clone TER-119) using MACS LS magnetic columns (Miltenyi Biotec). Generation of *Stat5a;Stat5b* double knockouts was carried out in cells from *Cas9; Bcl2* transgenic (tg) animals as described (Shin *et al*., 2021), or alternatively in cells from *Cas9*; *Bcl2*-tg; *Bcl11b^mCh/mCh^* mice in order to differentiate pre-commitment of BCL11B-nonexpressing cells from post-commitment BCL11B^+^ cells. For differentiation of DN cells in Artificial Thymic Organoid (ATO) cultures (**Figure S1A**), the protocol described by (Montel-Hagen *et al*., 2020) was followed, using stromal cells kindly supplied by A. Montel-Hagen and G. Crooks (UCLA). For differentiation in OP9-DLL1 cocultures (**Figure S1B-F**; **Figure S2**), enriched progenitor cells were co-cultured with OP9-DLL1 cells with IL-7 (Peprotech) and FLT3L (Peprotech) (10 ng/ml each) for 7 days in OP9 medium (α-MEM, 20% FBS, 2 mM glutamine, 100 IU/ml penicillin, 100 ug/ml streptomycin, and 50 µM β-ME). For RNA-Seq analysis (**Figure S1K and S1L and Table S3**), cells were collected and sorted to purify DN2 BCL11B^-^ and DN2 BCL11B^+^ subsets generated from both control and *Stat5a;Stat5b* double knockout cells with a *Bcl11b^mCh^* transgene. To generate DN2 cells for ChIP-Seq with and without IL-7 stimulation (**Figure S2**), precursors from conventional C57BL/6J mice were differentiated in OP9-DLL1 culture. For IL-7 stimulation, FACS-sorted DN2 (cKit^+^CD25^+^) cells that were deprived of IL-7 during the ∼6 hr purification were incubated with 20 ng/ml of IL-7 for 2 hours and compared with controls incubated without IL-7. All in vitro cultures were in a 37 °C, 7% CO_2_ environment.

To determine the cell surface protein expression by flow cytometry, cells were first incubated in 2.4G2 hybridoma cell supernatant to block Fc receptors. Cell surface staining was performed by incubating cells in a biotin-conjugated lineage cocktail (TCRβ, TCRγδ, CD19, NK1.1, CD11b, CD11c, and Ly6G/C) followed by secondary surface staining with fluorescently conjugated streptavidin, CD45, cKit, CD44, CD25, and hNGFR (for gRNA expressing vector). To exclude dead cells, a viability dye (Life Technologies, Aqua) or 7AAD (eBioscience) was used. To determine the STAT5 phosphorylation, cells were stimulated with IL-7 for 2 hr, fixed with 4% paraformaldehyde for 15 min at room temperature, and then permeabilized with 100% methanol at -20°C for 30 min prior to intracellular staining at room temperature for 45 min. Samples were acquired using a CytoFlex cytometer (Beckman Coulter), and data were analyzed with FlowJo version 10.8.1 (BD Biosciences). The following antibodies were used for flow cytometry analysis:

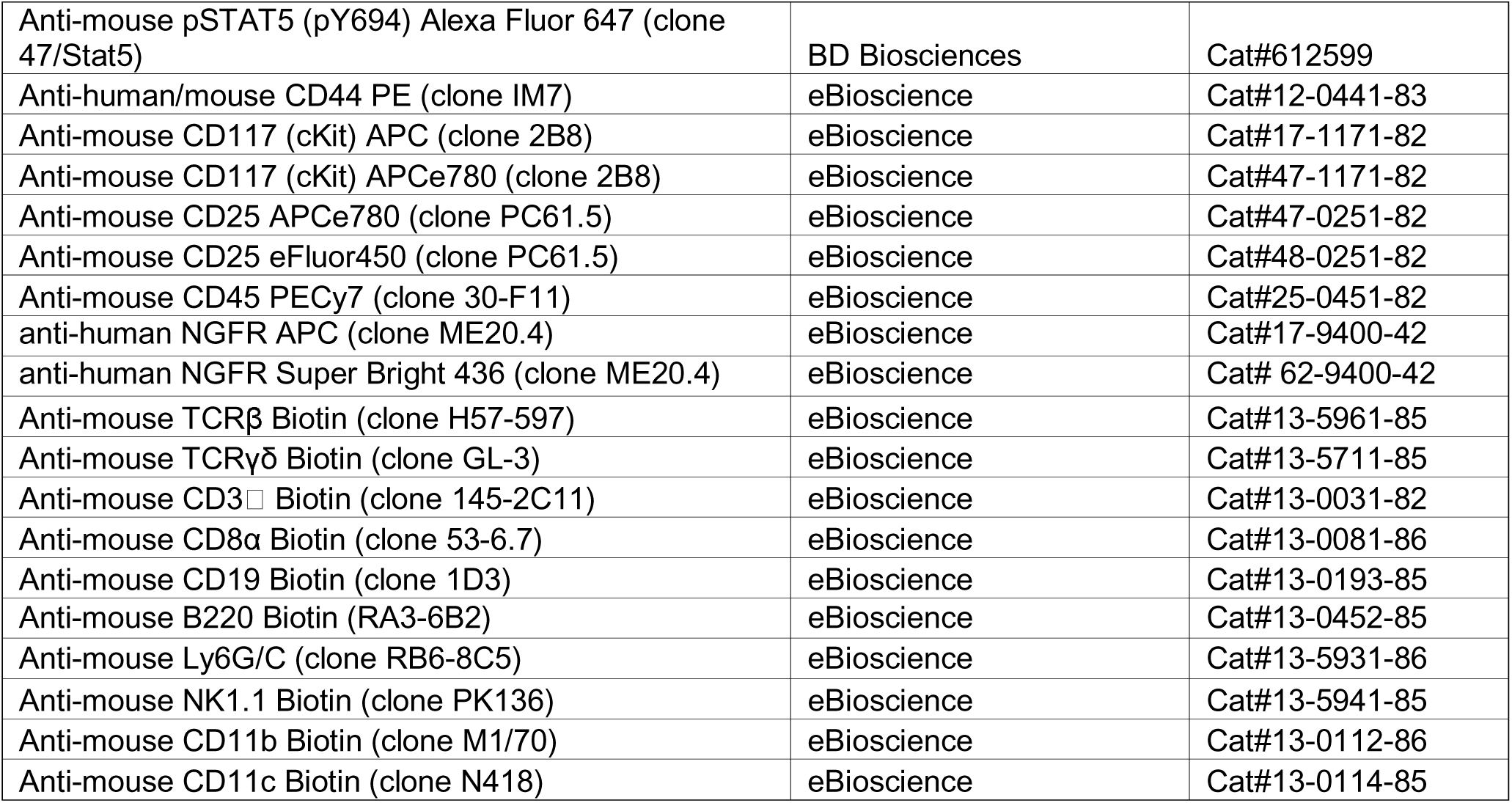

### Treg suppression assay

5 x 10^4^ Cell Trace Violet-labeled CD4^+^ T cells were cultured with irradiated APC (2000 rads) and soluble anti-CD3 (1 μg/ml) in the presence of purified splenic Tregs from either WT or CRISPR-Cas9 deleted mice. CD4^+^ T cells and splenic Tregs were purified using Stem Cell Technology bead-based kits. APC were prepared by depleting CD4^+^ and CD8^+^ T cells from spleen using Miltenyi Biotec magnetic beads. CD4^+^ T cells were labeled with Cell Trace Violet (Life Technologies) according to the manufacturer’s protocol. The CD4:Treg ratio of displayed flow cytometry profiles for CTV dilution was 2:1. APC were present in a 5-fold excess over CD4^+^ T cell numbers.

### Reporter assays

A DN3 cell line (SCID-ADH-2C2) that constitutively expresses CD25 was used. 5 x 10^6^ cells were electroporated with 2 μg of reporter plasmid containing constructs cloned 5’ of the minimal promoter of pNL3.1 and 0.2 μg pNL-TK in 20 μl buffer using Lonza P3 primary cell kit for 4D Nucleofector. Cells were immediately stimulated with cytokine for 24 hr and nanoluciferase activity was measured relative to control activity using the NanoGlo Dual Luciferase kit (Promega). The primers for cloning enhancer fragments into pNL3.1 are in **Table S3**.

### ChIP-Seq and RNA-Seq analysis

ChIP-Seq experiments were performed as previously described (Li *et al*., 2017; Lin et al., 2012) using the following antibodies: anti-STAT5B (Invitrogen 13-5300), anti-H3K27Ac (Diagenode), anti-MED1 (Bethyl A300-793A), anti-RAD21 (Abcam 217678), and anti-H3K4me1 (Abcam 176877). Libraries were prepared using the KAPA HyperPrep kit (Roche) and indexed with Illumina primers. PCR products were bar-coded (indexed) and sequenced using a HiSeq 2500 or NovaSeq6000 (both from Illumina). Sequenced reads (50 bp, single end) were obtained with the Illumina CASAVA pipeline and mapped to the mouse genome mm10 (GRCm38, Dec. 2011) using Bowtie 2.2.6 (Langmead et al., 2009) and Tophat 2.2.1 (Trapnell et al., 2009). Only uniquely mapped reads were retained. The mapped outputs were converted to browser-extensible data files, which were then converted to binary tiled data files (TDFs) using IGVTools 2.4.13 (Robinson et al., 2011) for viewing on the IGV browser (http://www.broadinstitute.org/igv/home). TDFs represent the average alignment or feature density for a specified window size across the genome. For ChIP-Seq data, we mapped reads into non-overlapping 20 bp windows for various transcription factors (STAT5B, MED1) and histone modifications H3K27ac. The reads were shifted 100 bp from their 5′ starts to represent the center of the DNA fragment associated with the reads. For RNA-Seq data, raw counts that fell on exons of each gene were calculated and normalized by using RPKM (Reads Per Kilobase per Million mapped reads). Differentially expressed genes were identified with the R Bioconductor package “edgeR” (Robinson et al., 2010). RNA-Seq analysis of in vitro-derived control and *Stat5a;Stat5b* double knockout DN2 cells was carried out using conditions described previously (Shin *et al*., 2021).

### HiChIP Protocol

HiChIP assays for H3K27Ac modification were performed using pre-activated CD8^+^ T cells from either wild-type or mutant mice, as described previously (Chandra et al., 2021). Briefly, cells were crosslinked with 1% formaldehyde and flash-frozen in liquid nitrogen. Fixed cells were lysed to obtain the nuclear fraction, and then chromatin was digested using intact nuclei and 200 U of the 4-base cutter Mbo I (New England Biolabs), and restricted ends religated as described Mumbach et al., 2017). Pelleted nuclei were dissolved in 130 μl of nuclear lysis buffer (50 mM Tris-HCl, pH 7.5, 10 mM EDTA, and 1% SDS) and sonicated using a Covaris S220 for 4 min. Sonicated chromatin was diluted ten times in ChIP Dilution Buffer and immunoprecipitation was done overnight at 4°C by incubating 2.5 μl (7 μg) of anti-H3K27ac antibody (C15410196; Diagenode) precoated on 25-μl protein A-coated magnetic beads (Thermo Fisher Scientific). Immunocomplexes were captured and washed three times for 5 min each with RIPA buffer, high-salt buffer, LiCl buffer and low-salt buffer (10 mM Tris-HCl, pH 8, 1 mM EDTA, and 50 mM NaCl). Beads were resuspended in TE (10 mM Tris-HCl, pH 8, 1 mM EDTA), transferred to fresh tubes, captured, and then resuspended in 200 μl of elution buffer (50 mM NaHCO3 and 1% SDS). Samples were treated with 10 μg of RNase A (Thermo Fisher Scientific) for 30 min at 37°C, then with 2 μl of 20 mg ml^−1^ proteinase K (Thermo Fisher Scientific) for 1 h at 55° C, and then incubated overnight at 65° C. DNA was purified using affinity columns (Zymo Research) and eluted in 20 μl of DNA elution buffer (10 mM Tris-HCl). Adapters were ligated to DNA using NEBNext Ultra DNA Library Prep Kit (New England Biolabs) according to the manufacturer’s protocol. Streptavidin C-1 beads were used to capture biotinylated DNA according to the manufacturer’s protocol and resuspended in 20 μl of DNA elution buffer. To generate the sequencing library, PCR amplifications of the DNA were performed while the DNA was still bound to the beads. Purified HiChIP libraries were size-selected to 300–800 bp using AMPure XP beads (Beckman Coulter Life Sciences) according to the manufacturer’s protocol and subjected to 2 × 50-bp paired-end sequencing on an HiSeq 2500 or NovaSeq6000.

### HiChIP Analysis

Reads were mapped as follows: For individual samples in different cell types, we applied the HiC-Pro pipeline on the respective paired-end reads. Each was mapped independently to the mouse mm10 reference genome, using the aligner bowtie2. Aligned reads were then paired, and paired reads involving two different Mbo I restrictions sites were retained. FitHiChIP (Bhattacharyya et al., 2019) was applied on the input set of valid HiChIP reads, and 1 kb binning was applied. Since the point of junction does not create a new Mbo I site, custom indexing is not required. FitHiChIP (Bhattacharyya *et al*., 2019) uses the valid read pairs generated by HiC-Pro (Servant et al., 2015) and a set of reference ChIP–Seq or HiChIP peaks corresponding to the target protein or histone modifications of interest (here H3K27ac) to derive statistically significant interactions. FitHiChIP applies fixed-size binning (here 5 kb) on the input set of valid HiChIP reads, and attributes a bin as peak-bin if that bin overlaps with a ChIP– Seq or HiChIP-inferred peak in the reference peaks file, subject to a 1-bp minimum overlap without any slack. Otherwise, the bin is labeled as a non-peak-bin. The background (set of locus pairs used to infer the null model) as well as the foreground (that is, set of locus pairs that were assigned a significance estimate) of FitHiChIP can be either peak-to-peak (that is, interactions between two peak bins) or peak-to-all (that is, interactions involving peak bins in at least one end). We used the default mode (peak-to-all pairs) for the foreground, which is the setting employed in this study. Note that FitHiChIP estimates the background contact probability and employs the binomial distribution on the generated contact probabilities to estimate P values, which are then corrected for multiple testing. Interactions having false discovery rate (FDR) < 0.01 are considered significant and reported as loop calls. These statistical analyses support the validity of interactions identified by FitHiChIP. Lack of loop counts (*Il2ra* gene promoter to regions close to UP1-6) in FitHiChIP-L (that detect peak-to-all interactions) in ΔUP1-6 KO cells also suggest that these are not from closely associated open chromatin regions. Loops calls were visualized using the Washington University Epigenome Browser [http://epigenomegateway.wustl.edu/browser/].

---

**Figure S1.**
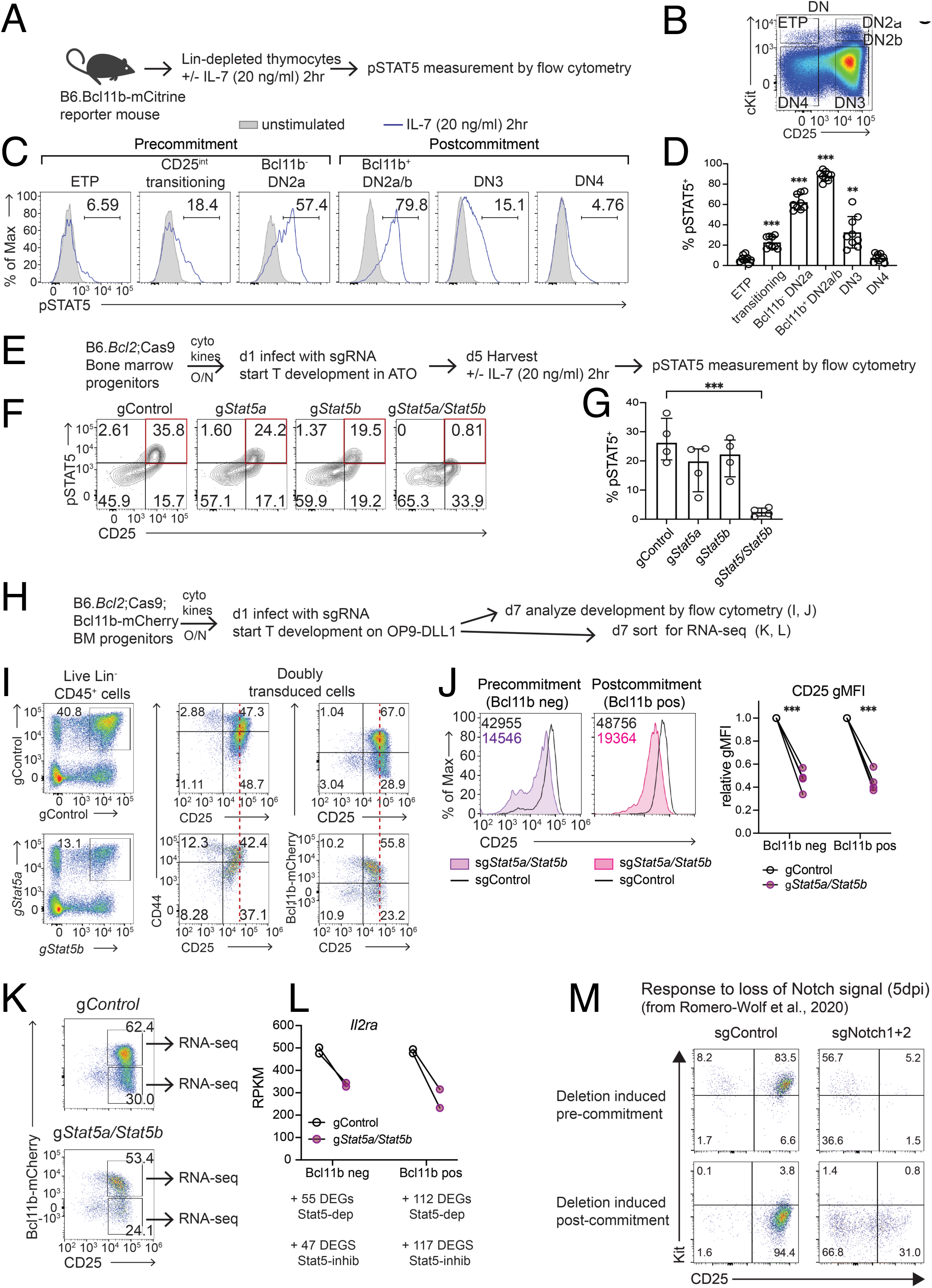
Regulation of *Il2ra* by STAT5A and STAT5B and Notch signaling in developing DN thymocytes. **(A-D)** Developmental time window for STAT5 activation by IL-7 in developing DN thymocytes. STAT5 phosphorylation profiles in different subsets of DN thymocytes in the acute response to IL-7 are shown. Thymocytes were from *Bcl11b-mCitrine* knock-in reporter mice (Kueh et al., 2016), which provide a marker sharply distinguishing newly committed from uncommitted cells within the DN2a stage. **(A)**Experimental scheme. **(B)** Flow cytometry gating strategy to define subsets of DN populations, based on expression of cKit, CD44, CD25, and the Bcl11b-mCitrine reporter. Mature cells or non-T-lineage cells in total thymocytes were first depleted (CD8α, TCRβ, TCRγδ, CD11b, CD11c, Ly6G/C, NK1.1, Ter119-depleted) to obtain thymic DN cells. **(C)** STAT5 phosphorylation patterns in different stages of DN cells in response to IL-7 stimulation. DN thymocytes were stimulated with IL-7 for 2 hrs in vitro, gated as shown in (B), and STAT5 phosphorylation measured by flow cytometry. Blue lines, with IL-7 stimulation; gray fill, unstimulated controls. **(D)**Histograms display frequency of pSTAT5^+^ cells in each subset over the background in unstimulated cells, n=9; **, p<0.01; ***, p<0.001, One-way ANOVA (in comparison to ETP). **(E-G)** Efficiency of acute Cas9-mediated deletion of both STAT5A and STAT5B in pro-T cells. **(E)** Experimental scheme. Progenitors were subjected to CRISPR disruption of *Stat5a* and/or *Stat5b* by introducing the indicated guide RNAs or controls, then induced to begin T cell development in artificial thymic organoid (ATO) culture *in vitro* (Montel-Hagen et al., 2020), an alternative to OP9-DLL1 culture. They were harvested at a timepoint when they were transitioning from the ETP to the DN2a and DN2b stages, which is expected to be the peak of STAT5 responsiveness (cf. **C-D**), and then tested for STAT5 phosphorylation in response to short-term IL-7 exposure. Note that the mouse genotype contains not only *Cas9* but also the *Bcl2* transgene to keep *Stat5*-deficient cells alive. ATO, artificial thymic organoid culture. **(F)** Ablation of IL-7-induced STAT5 phosphorylation when *Stat5a* and *Stat5b* are both deleted. Control confirms that the STAT5 response normally appears as cells activate CD25 expression in ETP to DN2a transition; this response was lost in *Stat5a, Stat5b* double knockout mice. **(G)** Summary of results from 4 independent experiments, *** *P* < 0.001, one-way ANOVA (in comparison to gControl). **(H-L)** Impact of *Stat5a/Stat5b* double deletion on natural developmentally induced *Il2ra* (CD25) expression in pro-T cells. **(H)** Schematic of experiment. Control guide RNAs or guide RNAs to induce *Stat5a/Stat5b* deletion were introduced just before the cells started T-cell differentiation. The time of harvest was chosen to enable generation of both pre-and post-commitment CD25^+^ (*Il2ra-*expressing) cells, corresponding to DN2a, DN2b, and some DN3 cells. Both *Bcl2* and *Cas9* transgenes were included as in (E). The *Bcl11b-mCherry* transgene, like the *Bcl11b-mCitrine* transgene in (A-D), provides a sensitive indicator of the DN2-DN3 cells’ transition through commitment. **(I)** Left, gating of cells transduced with control guide RNA constructs (sgControl) or guide RNAs against *Stat5a* and/or *Stat5b*; right, expression of CD44, CD25 and Bcl11b-mCherry (commitment marker) in the *Stat5a*/*Stat5b* doubly-transduced cells. Vertical broken line assists comparison of CD25 expression levels. **(J)**Histograms of CD25 expression in pre-commitment (left) and post-commitment (right) DN2-DN3 cells from control and *Stat5a*/*Stat5b* double knockout mice. Numbers in the upper left denote geometric Mean Fluorescent Intensities (gMFI). Right, Changes in CD25 gMFI in *Stat5a*/*Stat5b*-deficient cells relative to controls in independent experiments, n=4, \*\*\**P* < 0.001, paired t-test. **(K)** Developing pro-T cells were sorted as shown (left) to collect samples for RNA-Seq analysis. In addition to CD25 (encoded by *Il2ra*) as a developmental stage marker, *Bcl11b-mCherry* was used as an independent marker to monitor the transition through commitment, which normally occurs after CD25 upregulation. **(L)** Expression of transcripts (RPKM) from the *Il2ra* locus in control and *Stat5a*/*Stat5b* knockout DN2-DN3 cells, before (left) and after (right) commitment, in two independent experiments. Below, numbers of genes from whole RNA-Seq datasets that reproducibly scored as differentially expressed genes (DEGs) in *Stat5a/Stat5b* double knockout cells as compared to controls are given separately for the Bcl11b reporter-negative cells (before commitment) and for the Bcl11b reporter-expressing cells (after commitment). Criteria for DEGs: genes with average FPKM > 1 and adjusted p value <0.05; this criterion also excluded genes with log_2_ fold change < 1 in either direction. Full details of DEGs and their linkage to STAT5B binding sites are presented in Table S3. Stat5-dep, higher in control than in *Stat5a, Stat5b* double knockout; Stat5-inhib, higher in double knockout than in control. (**M**) Regulation of *Il2ra* by Notch signaling in DN2-3 pro-T cells. These data were reproduced from a previous report (Romero-Wolf *et al*., 2020) showing the impact of acute disruption of *Notch1* and *Notch2* together, in pre-commitment (top) or post-commitment (bottom) pro-T cells. For deletion specifically in post-commitment cells, the *Cas9;Bcl2* transgenic cells were first cultured through commitment before adding the sgRNAs against *Notch1* and *Notch2* or control targets. Both pre- and post-commitment samples were harvested for analysis 5 days after sgRNA introduction. When Notch is removed, CD25 expression is downregulated in both cKit^+^ pre-commitment and cKit^-^ post-commitment cells.

**Figure S2:**
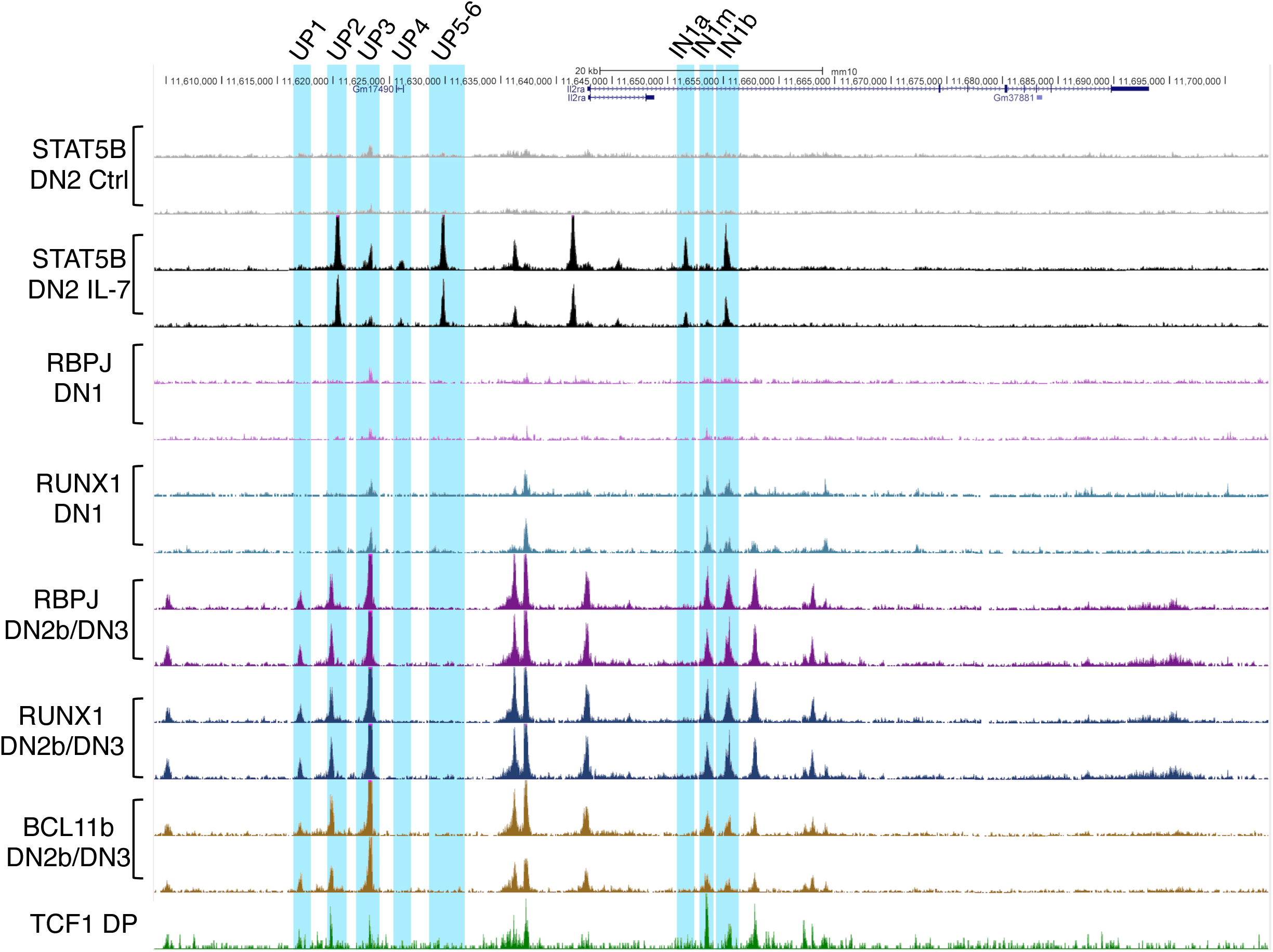
STAT5 binding sites are hubs of stage-specific, multi-transcription factor interaction. The genomic region of *Il2ra* is shown (chr2:11,603,933-11,703,852); the STAT5 binding elements as defined in **Figure 1** are highlighted in cyan. Tracks show ChIP-Seq data for STAT5B in two independent replicates each for control (IL-7 starved DN2 cells) and for DN2 cells treated with IL-7. These patterns are compared with published ChIP-Seq datasets showing the patterns of occupancy of RUNX1 and RBPJ (a NOTCH partner) in DN1 cells, before IL-7R is expressed, and for RBPJ, RUNX1, and BCL11b in DN2b/DN3 cells, after commitment. Independent biological replicates are shown for each. Also shown are binding data for TCF1 in DP thymocytes. The previously published track data are from accession numbers GSE110020, GSE148441, GSE110305, and GSE46662.

**Figure S3.**
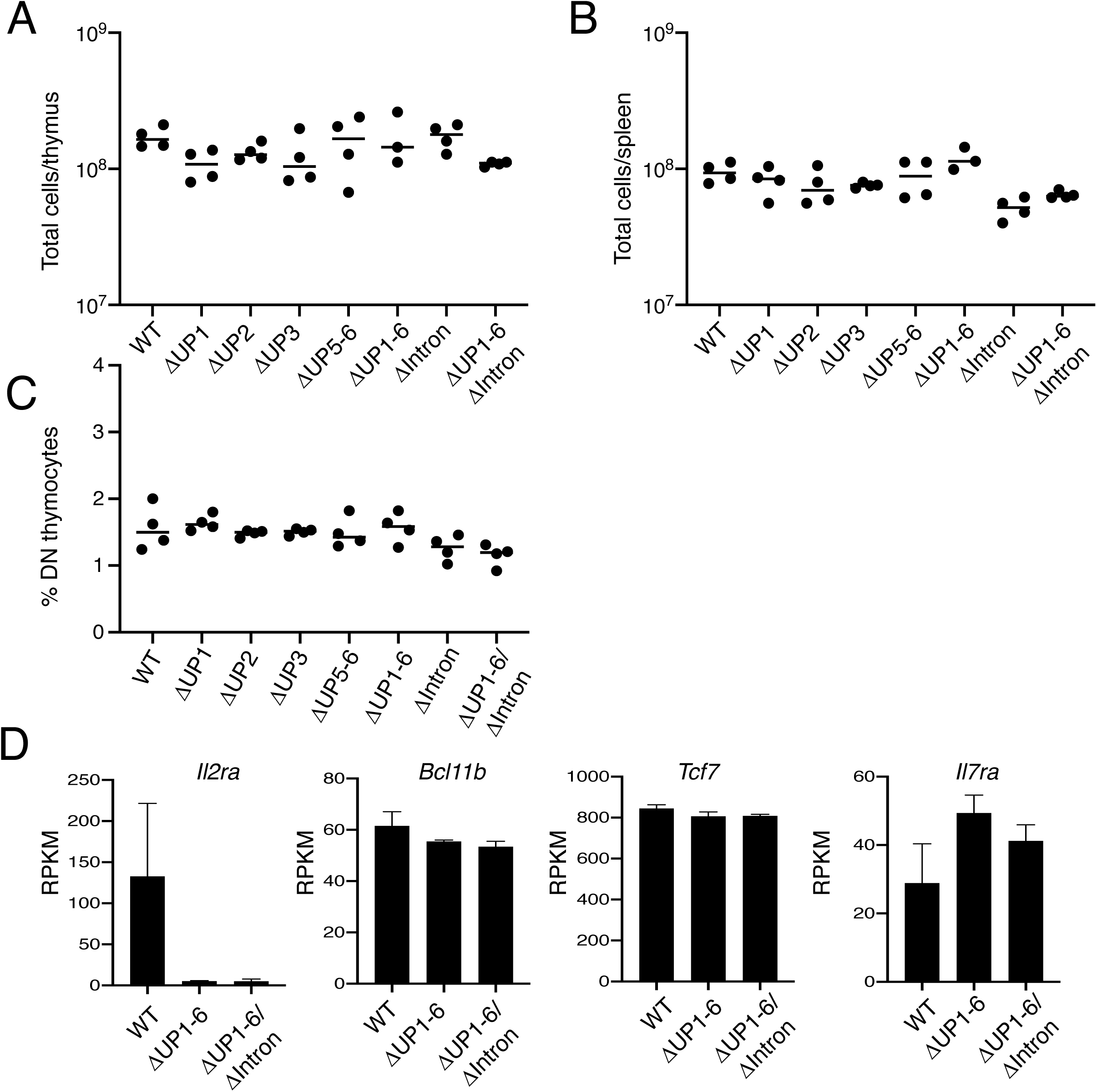
**(A and B)** Total thymic (**A**) and splenic (**B**) cellularity in WT and the indicated mutant mice. **(C)** Percentage of DN thymocytes relative to the total number of thymocytes. (**D**) RNA amounts as assessed by RNA-Seq in double negative thymocytes from WT, ΔUP1-6, and ΔUP1-6/ΔIntron mice. Average RPKM from two libraries is shown. FoxP3^+^ Tregs as a percentage of total CD4^+^ T cells in the thymus **(A)** and spleen **(B)**.

**Figure S4.**
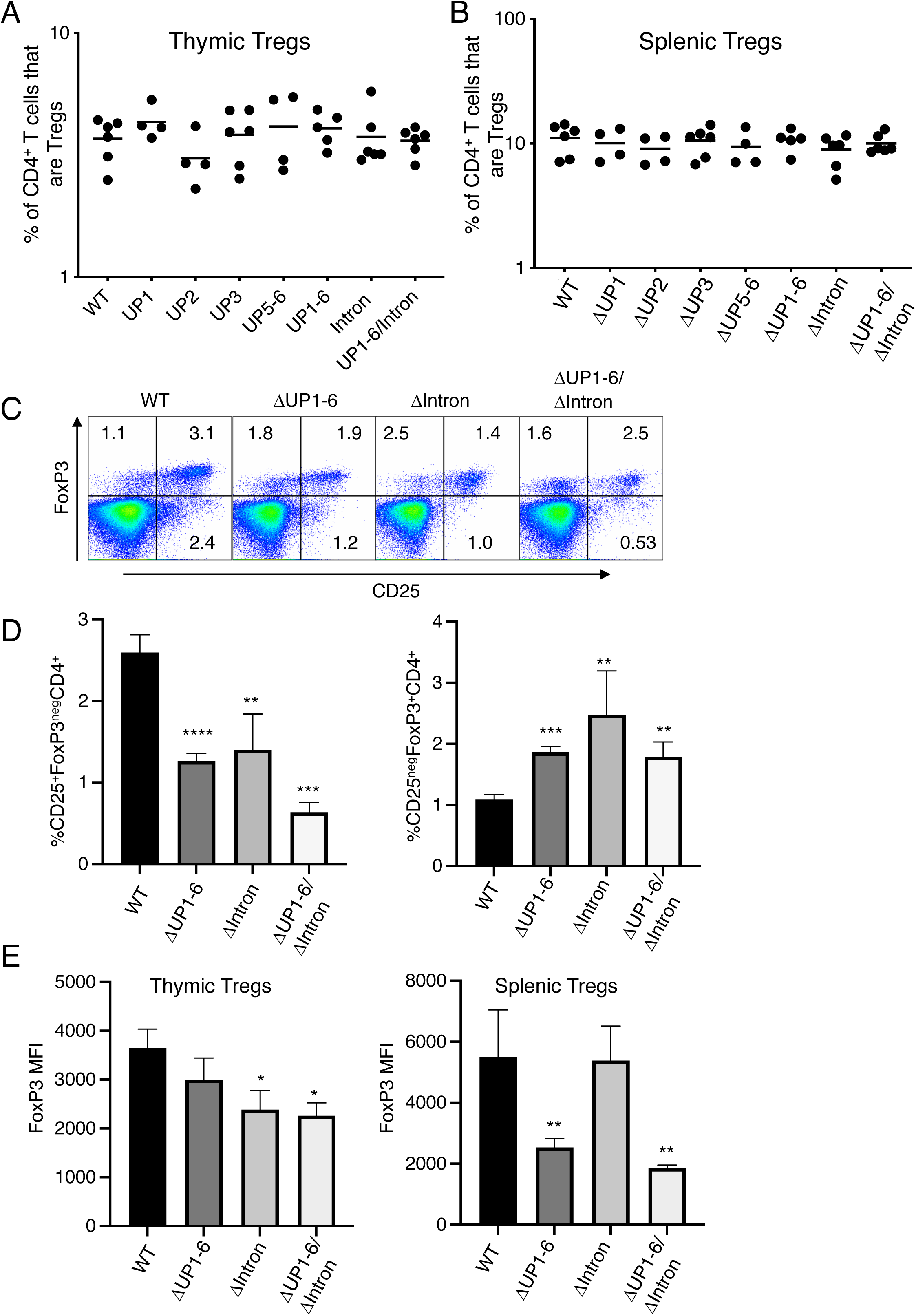
Flow cytometric analysis of Treg populations in the thymus. (**A and B**) FoxP3^+^ CD25^+^ Tregs as a percentage of total CD4^+^ T cells in the thymus **(A)** and spleen **(B). (C and D)** CD25 and FoxP3 expression were measured on CD4^+^ single positive thymocytes. Representative flow plots are shown in **(C)** and the percentage of CD25^+^FoxP3^neg^ or CD25^neg^FoxP3^+^ CD4^+^ cells is quantitated from in **(D)**. **(E)** FoxP3 MFI was measured in gated CD4^+^CD25^+^FoxP3^+^ Tregs in either thymus (left panel) or spleen (right panel). Data in **(D)** and **(E)** were accumulated from 3 experiments.

**Figure S5.**
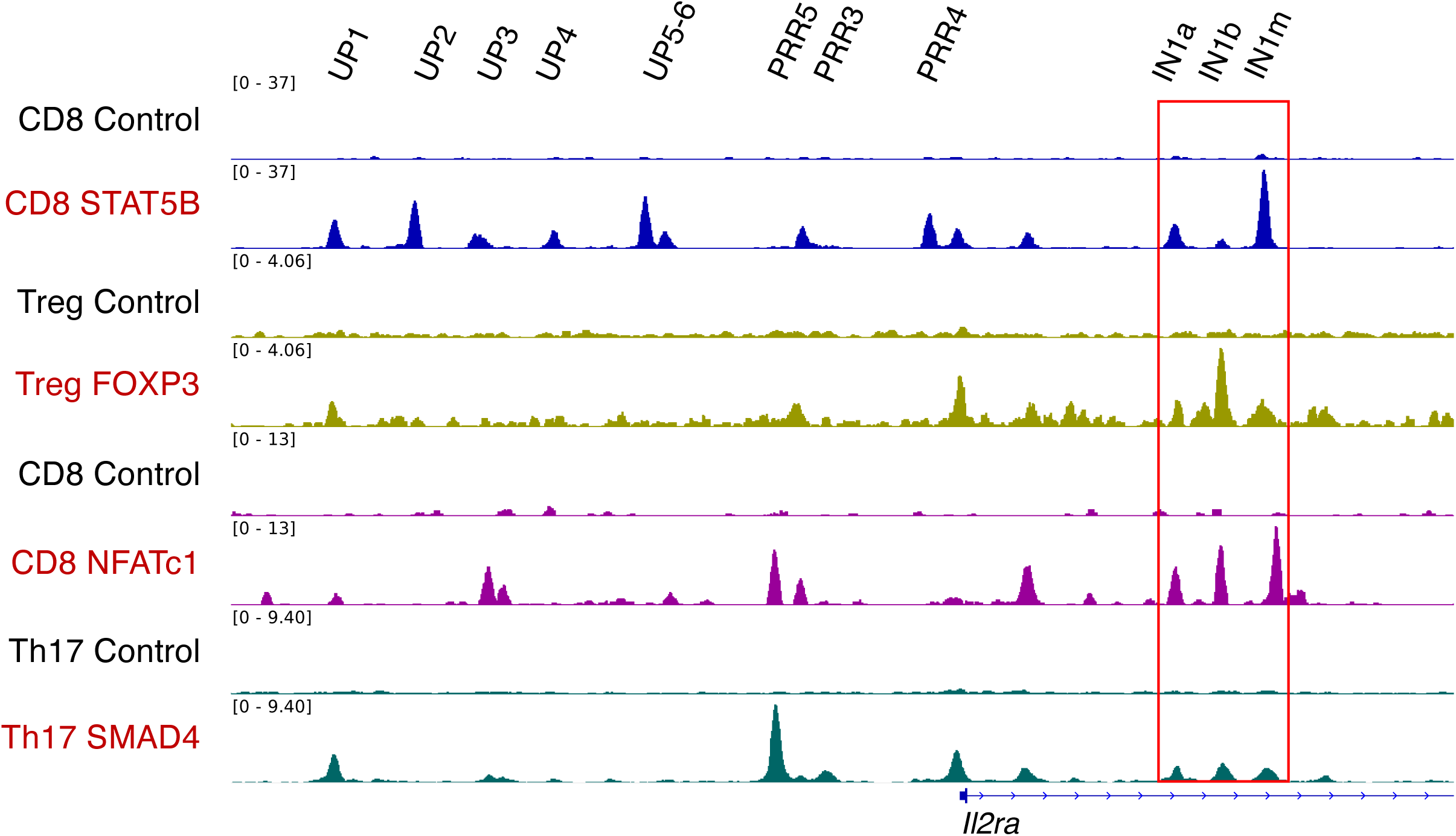
Alignment of binding profiles of STAT5B, NFATc1, FOXP3, TCF1 and SMAD4 at the *Il2ra* locus. IGV browser tracks showed the alignment of ChIP-Seq data in IL-2-activated STAT5 in CD8^+^ T cells, NFATc1 in CD8^+^ T cells, FOXP3 in Tregs, TCF1 in thymocytes, and SMAD4 in Th17 cells.

**Figure S6.**
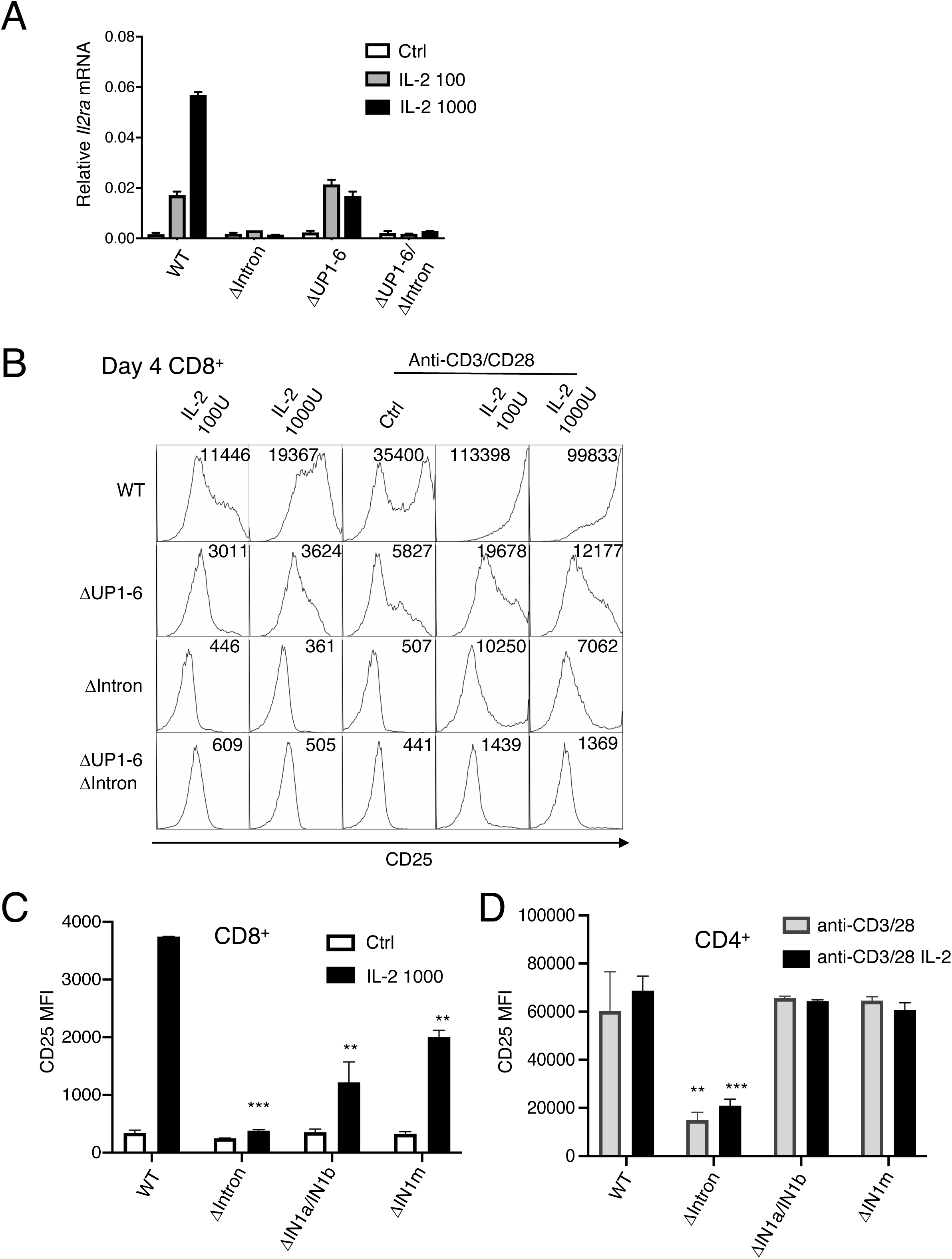
**(A)** RT-PCR was used to quantitate *Il2ra* mRNA expression in cells from WT, ΔIntron, or ΔUP1-6 CD8^+^ T cells stimulated for 2 days with IL-2 (100 or 1000 U/ml). **(B)** Flow cytometric measurement of CD25 in CD8^+^ T cells from WT, ΔIntron, ΔUP1-6, and ΔUP1-6/ΔIntron mice cultured for 4 days with or without IL-2 or IL-2+ anti-CD3 + anti-CD28. **(C)** Flow cytometric measurement of CD25 (MFI) in CD8^+^ T cells from WT, ΔIntron, ΔIN1a and IN1b, and ΔIN1m mice. CD8^+^ T cells were stimulated with or without IL-2 (1000 U/ml) for 3 days. (**D**) Flow cytometric measurement of CD25 MFI in CD4^+^ T cells stimulated with anti-CD3 + anti-CD28 in the presence or absence of IL-2 (1000 U/ml) for 3 days.

**Figure S7.**
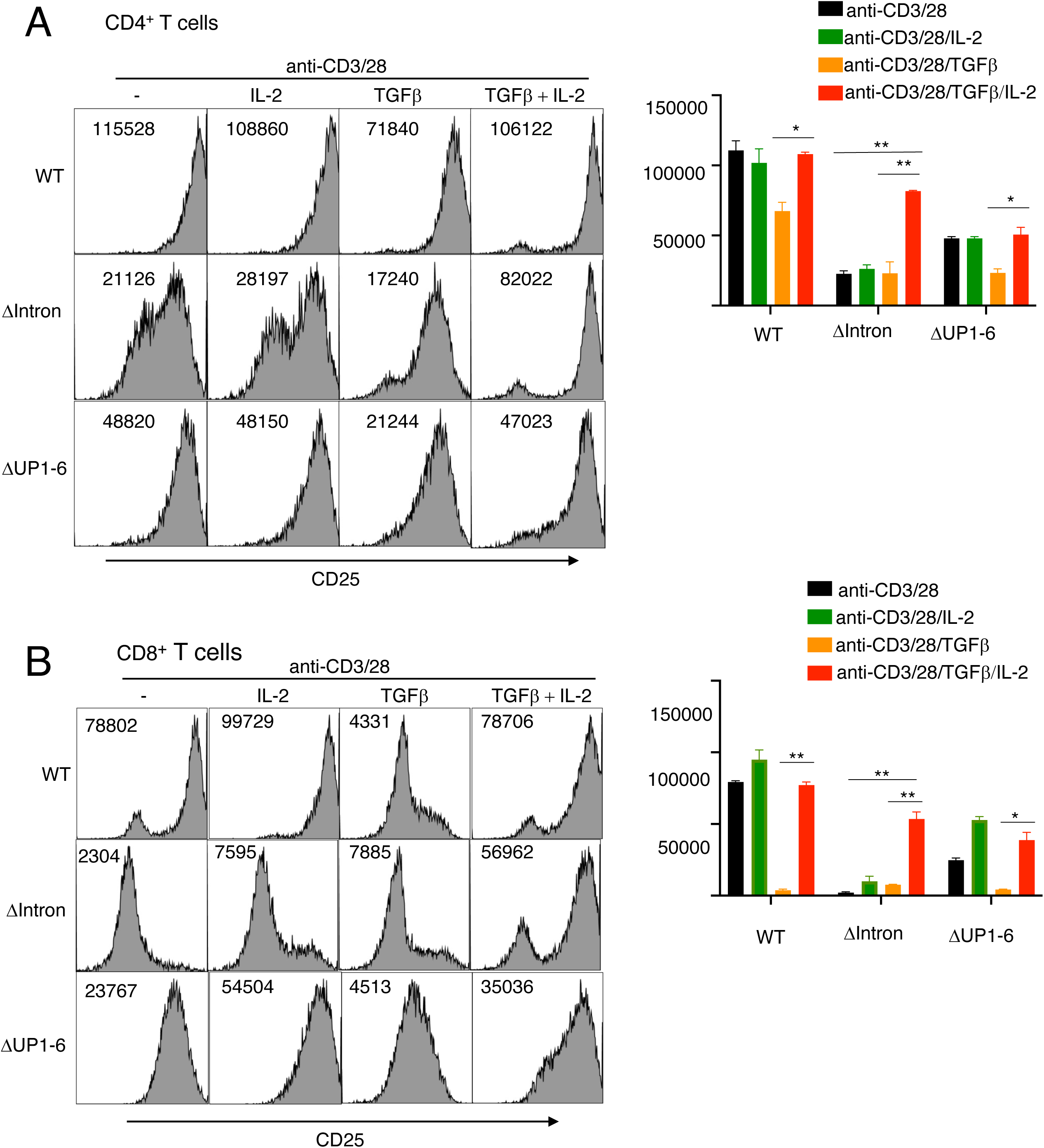
Effects of TGFβ on IL-2 induction of CD25 in CD4^+^ and CD8^+^ T cells. **(A)**CD4^+^ T or **(B)** CD8^+^ T cells were stimulated with anti-CD3 + anti-CD28 in the presence or absence of IL-2 either without TGFβ (left 2 panels) or in the presence of TGFβ (right 2 panels) for 3 days. CD25 was measured by flow cytometry, representative flow panels are shown on the left and quantitation is shown on the right.

**Figure S8.**
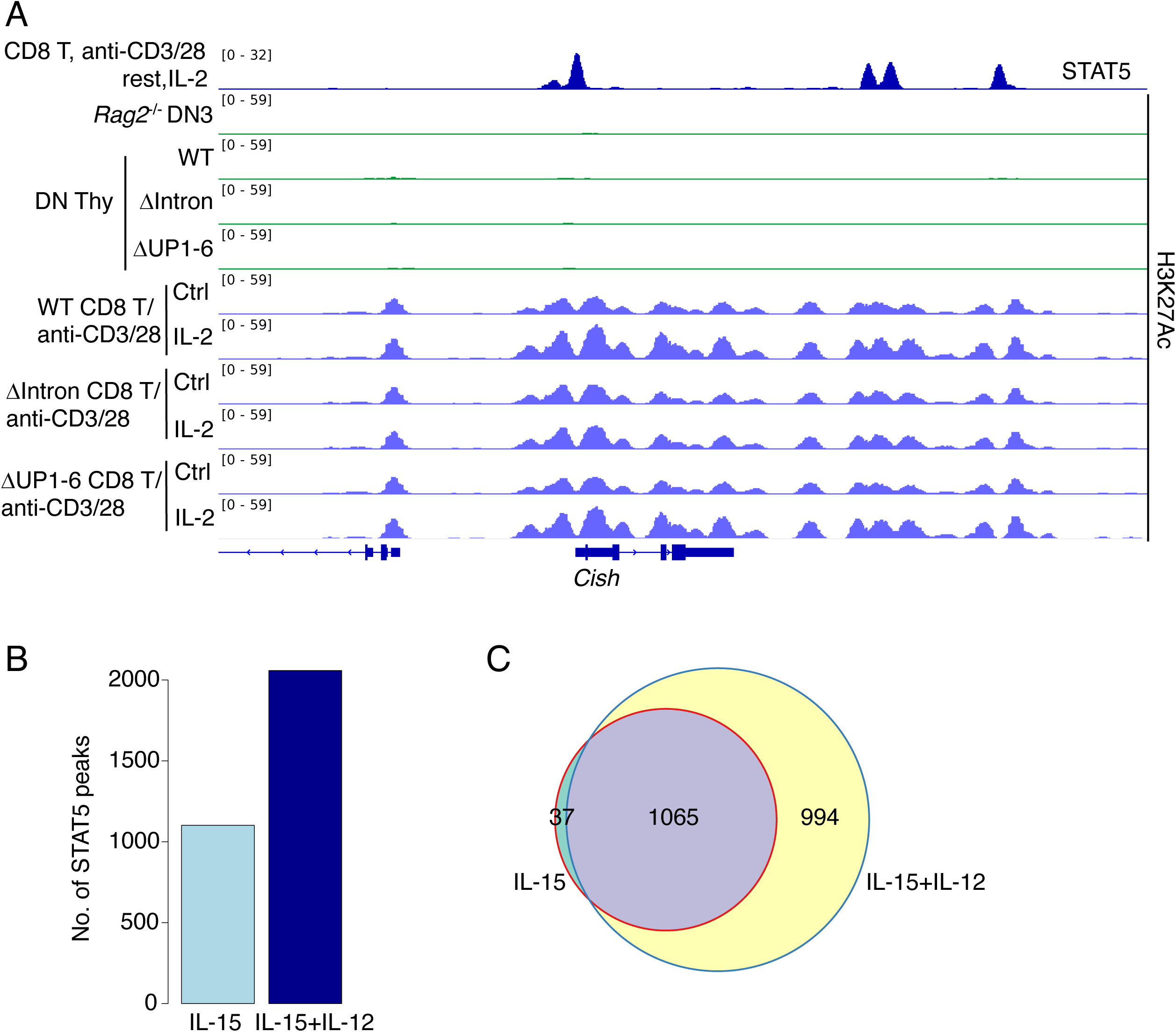
**(A)** H3K27Ac ChIP-Seq analysis at the *Cish* locus as a control for the effects at the *Il2ra* locus (shown in **Figure 6A**). H3K27Ac ChIP-Seq experiments were performed on two independent samples, with similar results. Representative data are shown. (**B and C**) The number of identified STAT5 binding sites (**B**) and Venn-diagram of shared STAT5 (**C**) sites in NK cells, treated with either IL-15 alone, or IL-15 + IL-12 are shown.

**Figure S9.**
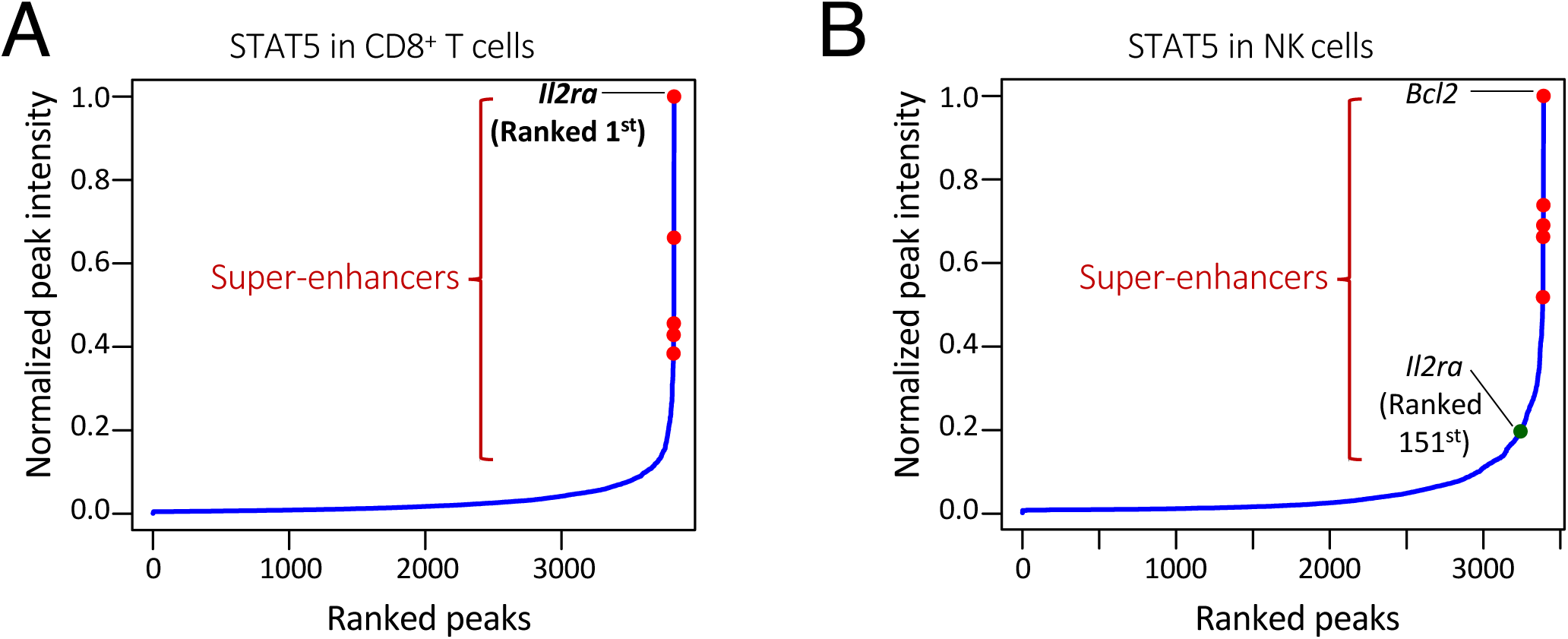
STAT5-bound super-enhancers in CD8^+^ T cells and NK cells. Super-enhancer analysis using IL-2-activated STAT5 in preactivated CD8^+^ T cells **(A)** and IL-15-activated STAT5 in NK cells **(B)**is illustrated. y axis, normalized density (also known as the super-enhancer score) of STAT5 binding; x axis, ranking of peaks from the lowest to highest super-enhancer score. Only protein-coding genes were analyzed and plotted.

**Figure S10.**
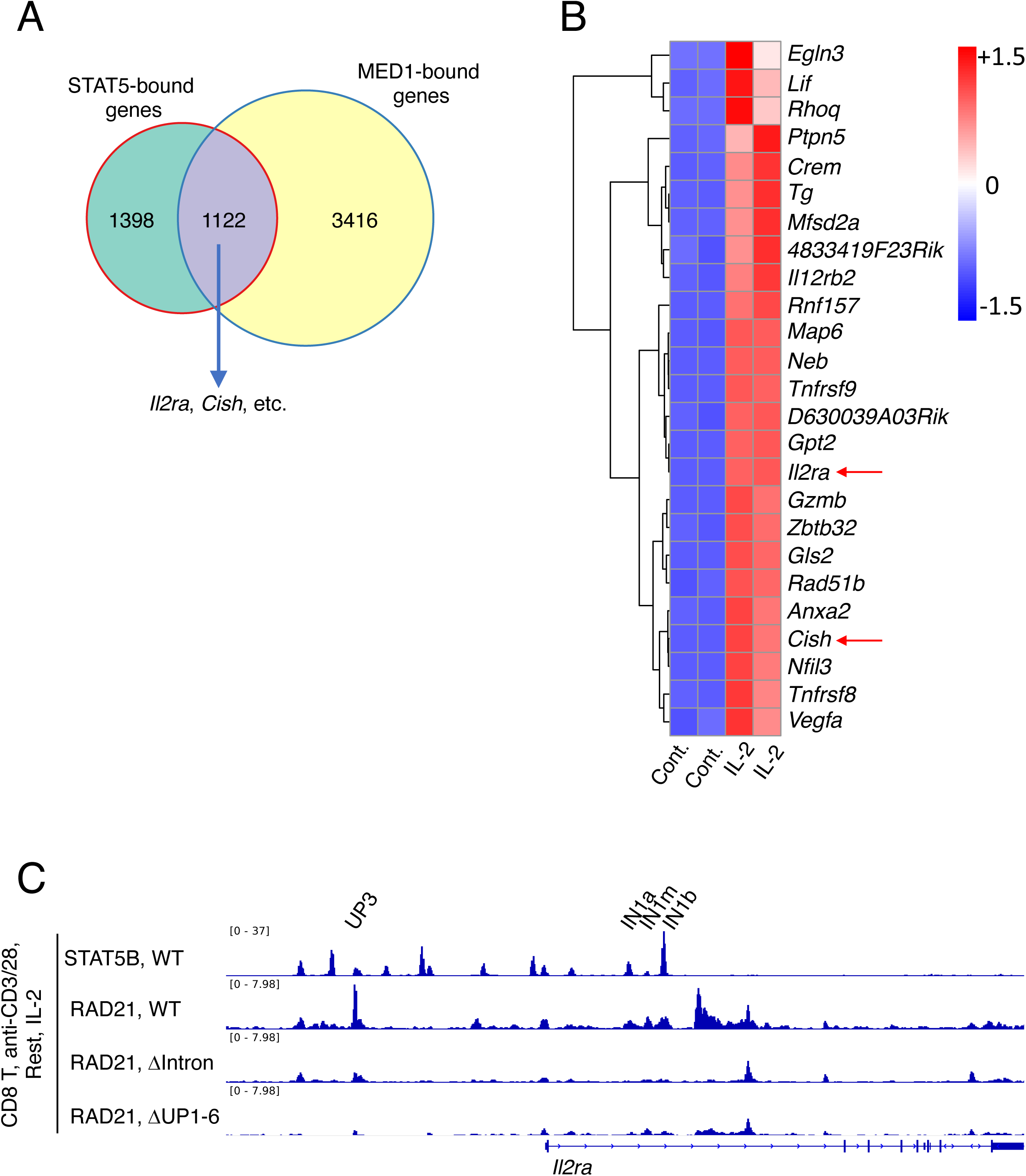
STAT5-bound genes were largely co-bound by MED1 (Mediator Complex Subunit 1). **(A)**Venn diagram showing 44.5% of genes bound by STAT5 were co-bound by MED1, including classical IL-2-inducible super-enhancer-containing genes, such as *Il2ra* and *Cish*. **(B)** Heat map reveals STAT5/MED1 co-bound genes were among the most potently IL-2-inducible genes (25 out of top 100, sorted by log_2_FC) in anti-CD3 + anti-CD28 pre-activated CD8^+^ T cells. **(C)** The binding of cohesin subunit RAD21 binding was determined by ChIP-Seq analysis; shown at the *Il2ra* locus. CD8^+^ T cells were stimulated with IL-2 prior to ChIP-Seq with anti-STAT5 or with anti-CD3 + anti-CD28 + IL-2 prior to ChIP-Seq with anti-RAD21. Experiments were performed on two independent samples, with similar results. Representative data are shown.

**Table S1.**
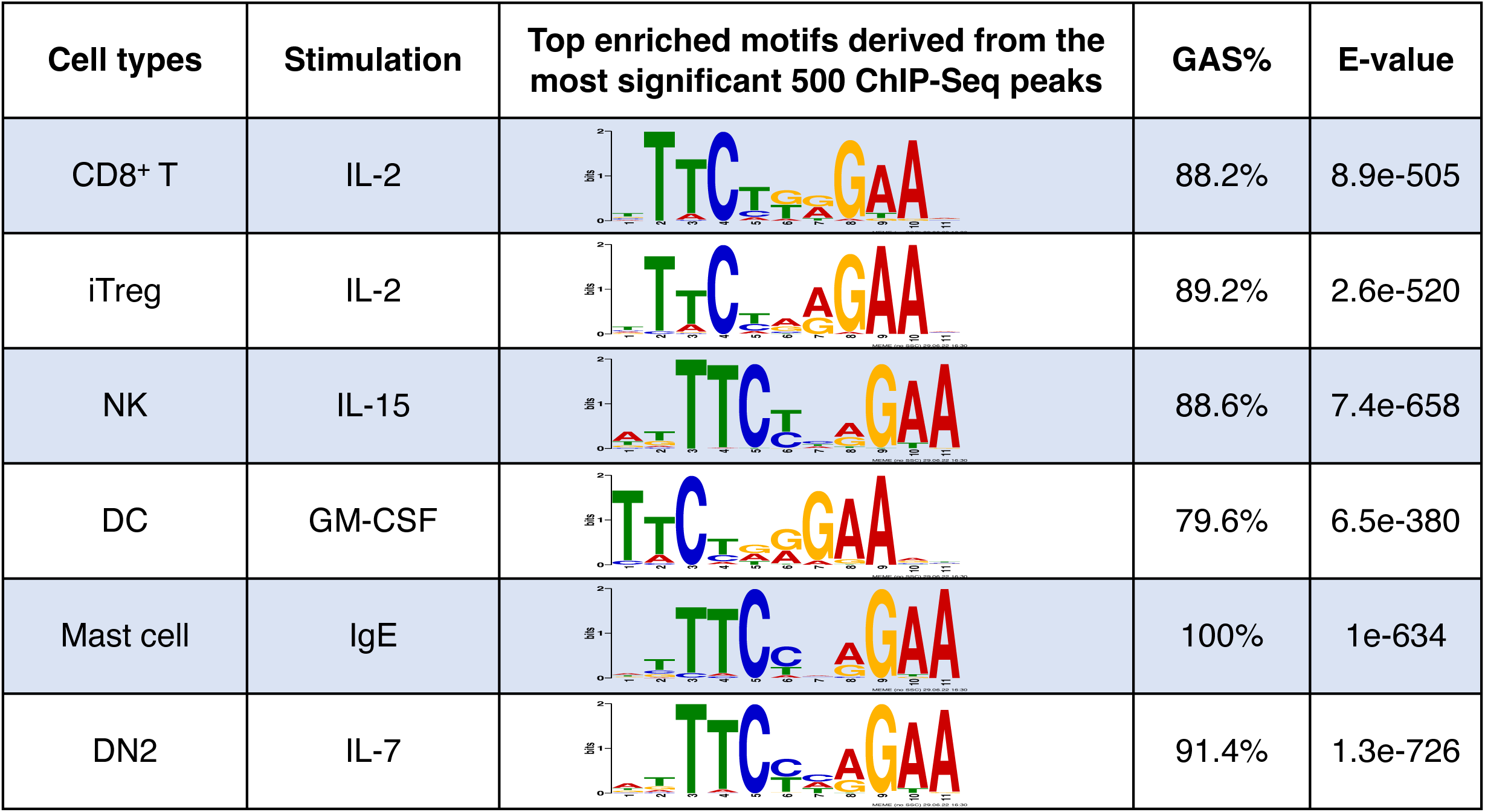
De novo motif analysis of STAT5 ChIP-Seq data in various cell types. For each STAT5 ChIP-Seq library in the indicated cell types, we identified binding sites (peaks) by comparing to the IgG control. We then selected the top 500 peaks with lowest FDR values, extracted 100 bp of DNA sequence centered on the peak summits, and performed de novo motif analysis using MEME to characterize the STAT5 consensus binding motifs in CD8^+^ T (IL-2), DN2 (IL-7), iTreg (IL-2), NK (IL-15), DC (GM-CSF) and mast cells (IgE). The most significant motifs are shown, as well as motif frequencies (GAS%) and the corresponding consensus E-values.

| Cell types | Stimulation | Top enriched motifs derived from the most significant 500 ChIP-Seq peaks | GAS% | E-value |
| --- | --- | --- | --- | --- |
| CD8<sup>+</sup> T | IL-2 |  | 88.2% | 8.9e-505 |
| iTreg | IL-2 |  | 89.2% | 2.6e-520 |
| NK | IL-15 |  | 88.6% | 7.4e-658 |
| DC | GM-CSF |  | 79.6% | 6.5e-380 |
| Mast cell | IgE |  | 100% | 1e-634 |
| DN2 | IL-7 |  | 91.4% | 1.3e-726 |

**Table S2.** STAT5 binding GAS motif at each Il2ra enhancer element. For each STAT5 binding site (peak), canonical GAS or GAS-like motifs with only single mismatches were shown with their chromosomal positions, E-values, and motif DNA sequence. MAST (Motif Alignment and Search Tool) was used to scan the GAS motif at each enhancer element.

| STAT5 peaks | chrom | start | end | Peak Score<br>-10log(pval) | GAS motif | E-value | Motif DNA<br>sequence |
| --- | --- | --- | --- | --- | --- | --- | --- |
| UP1 | chr2 | 11617088 | 11617096 | 961.8 | GASn | 7.45E-05 | TTCGAGGGA |
| UP2 | chr2 | 11620444 | 11620452 | 2017.56 | GASc | 0.00002 | TTCAATGAA |
| UP3 | chr2 | 11622823 | 11622831 | 475.59 | GASc | 1.63E-05 | TTCAGAGAA |
| UP4 | chr2 | 11626094 | 11626102 | 775.34 | GASc | 1.79E-06 | TTCTAAGAA |
| UP5 | chr2 | 11629859 | 11629867 | 2032.24 | GASc | 4.63E-06 | TTCTGAGAA |
| UP6 | chr2 | 11630596 | 11630604 | 301.23 | GASn | 1.16E-04 | TCCACAGAA |
| UP7/PRR5 | chr2 | 11636314 | 11636322 | 570.14 | GASc | 5.47E-06 | TTCACAGAA |
| PRR3 | chr2 | 11641489 | 11641497 | 700.8 | GASc | 2.36E-06 | TTCTGAGAA |
| Promoter | chr2 | 11642554 | 11642562 | 533.34 | GASn | 3.86E-05 | TGCCAAGAA |
| PRR4 | chr2 | 11645576 | 11645584 | 340.29 | GASc | 6.64E-06 | TTCTAGGAA |
| IN1a | chr2 | 11651616 | 11651624 | 1053.78 | GASc | 7.68E-07 | TTCAAAGAA |
| IN1m | chr2 | 11653492 | 11653500 | 172.97 | GASn | 4.04E-05 | TTCCAGGGA |
| IN1b | chr2 | 11655221 | 11655229 | 2802.08 | GASc | 3.04E-05 | TTCTCAGAA |

**Table S3.** STAT5 responsive genes in BCL11b^-^ and BCL11b^+^ DN2 cells upon acute deletion of *Stat5a* and *Stat5b* using CRISPR/Cas9. Legend and methodological features of the analysis are given on the “Readme” page.

